# Microvascular engineering for the development of a non-embedded liver sinusoid with a lumen: when endothelial cells do not lose their edge

**DOI:** 10.1101/2023.10.31.564881

**Authors:** Ana Ximena Monroy-Romero, Brenda Nieto-Rivera, Wenjin Xiao, Mathieu Hautefeuille

## Abstract

Microvascular engineering seeks to exploit known cell-cell and cell-matrix interactions in the context of vasculogenesis to restore homeostatic or disease development of reliable capillary models *in vitro*. However, current systems generally focus on recapitulating microvessels embedded in thick gels of extracellular matrix, overlooking the significance of discontinuous capillaries, which play a vital role in tissue-blood exchanges particularly in organs like the liver. In this work, we introduce a novel method to stimulate the spontaneous organization of endothelial cells into non-embedded microvessels. By creating an anisotropic micropattern at the edge of a development-like matrix dome using Marangoni-flow, we achieved a long, non-random orientation of endothelial cells, laying a premise for stable lumenized microvessels. Our findings revealed a distinctive morphogenetic process leading to mature lumenized capillaries, demonstrated with both murine and human immortalized liver sinusoidal endothelial cell lines (LSECs). The progression of cell migration, proliferation and polarization was clearly guided by the pattern, initiating the formation of a multicellular cord that caused a deformation spanning extensive regions and generated a wave-like folding of the gel, hinged at a laminin depleted zone, enveloping the cord with gel proteins. This event marked the onset of lumenogenesis, regulated by the gradual apico-basal polarization of the wrapped cells, leading to the maturation of vessel tight junctions, matrix remodeling, and ultimately the formation of a lumen—recapitulating the development of vessels *in vivo*. Furthermore, we demonstrate that the process strongly relies on the initial gel edge topography, while the geometry of the vessels can be tuned, from a curved to a straight structure. We believe our facile engineering method, guiding an autonomous self-organization of vessels without the need for supporting cells or complex prefabricated scaffolds, holds promise for future integration into microphysiological systems featuring discontinuous, fenestrated capillaries.

**Graphical abstract:** 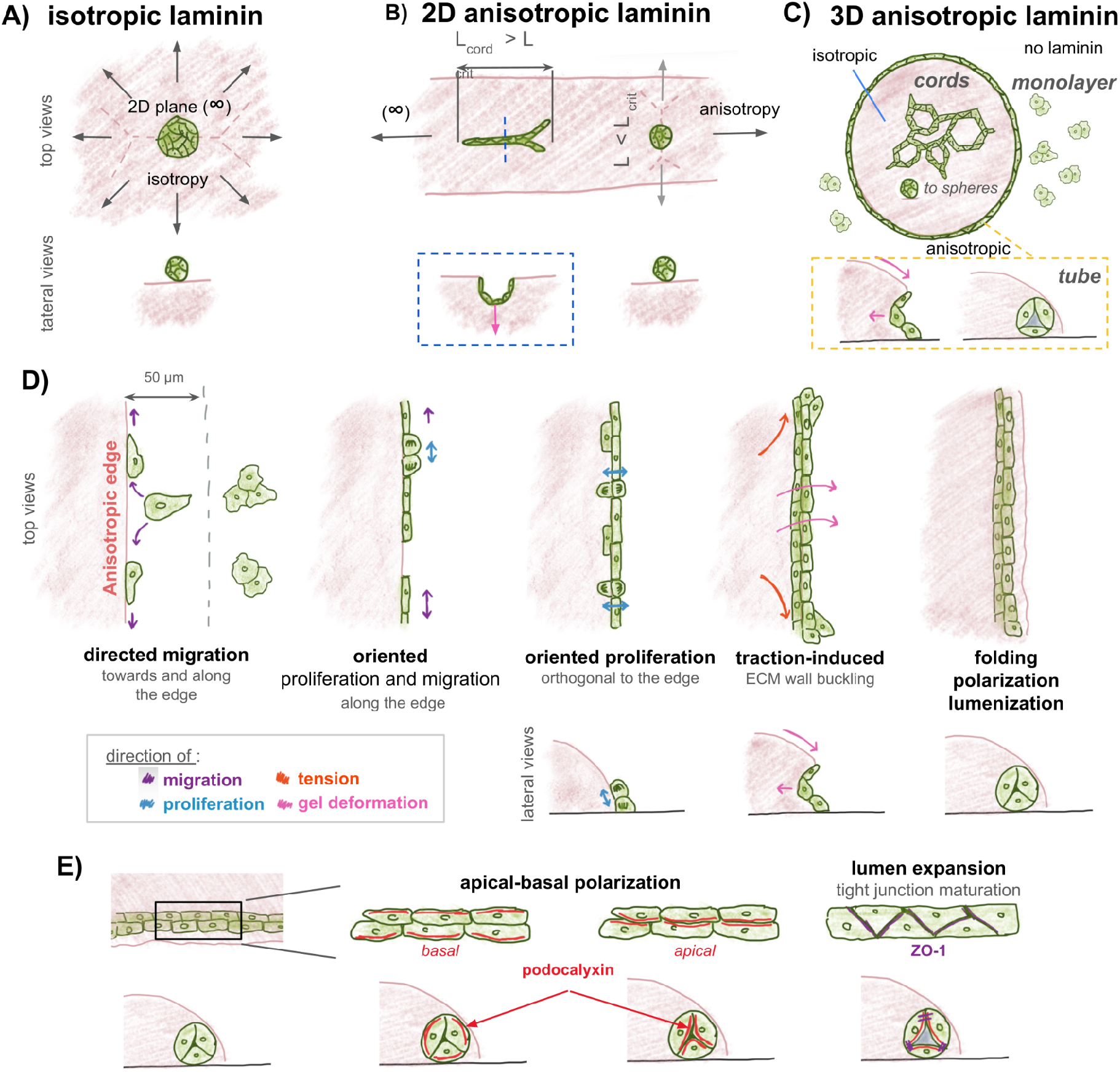

## 1. Introduction

The development of perfusable tubular structures that closely represent the physiological architectures of blood vessels is of growing interest ^1,2^. Recently, it has been proposed that vascular engineering could benefit from new discoveries in cell biology to harness developmental processes for bottom-up engineering of functional vessels ^3^. These processes include the *de novo* vessel formation or vasculogenesis and the sprouting of new vessels from pre-existing ones or angiogenesis ^4^. Using biomaterials and microfabrication techniques, tissue engineering at the microvessel-level can be precisely controlled to study and guide tissue morphogenesis^5^. For example, regarding vasculogenesis, disaggregated human umbilical vein endothelial cells (HUVEC) have been embedded in a fibrin hydrogel crosslinked inside a microfluidic chip. Under static conditions or controlled interstitial flow, these cells formed stable lumenized capillaries that then were directly perfused inside the chip from the lateral channels ^6^. Alternatively, regarding angiogenesis, a microfluidic channel incorporating a porous collagen I hydrogel has been used to generate gradients of vascular endothelial growth factor (VEGF), which guided angiogenic sprouts from side channels hosting HUVEC monolayers until anastomosis was achieved and the vessels were perfused ^7^. Both of these experimental setups allow for co-culture with support cells like fibroblasts, pericytes and immune cells to improve tissue recapitulation with vascularisation. Furthermore, they have successfully recreated vascularized tumor microenvironments ^8^, promising great progress in vascular engineering applications.

However, in all these solutions, the engineering strategy results in embedded vessels within extracellular matrix (ECM) gels, whose thickness can vary between hundreds of micrometers to one millimeter, depending on the fabrication process. This encapsulation specificity is unfortunately incompatible when seeking to imitate specialized capillaries such as the liver sinusoids. These vessels are highly permeable and possess a discontinuous basement membrane (BM) mainly composed of laminin and collagen IV ^9^. Liver sinusoidal endothelial cells (LSECs), which compose these capillaries, feature microscopic *fenestrae* on their surface ^10^, allowing for controlled passage of metabolites. They also express various mechanosensors which make them highly responsive to pressure, shear and changes in surrounding stiffness ^11,12^ which can in turn significantly impact the entire microenvironment in both homeostasis preservation and disease development ^13,14^. Still, the mechanisms at play remain unclear, primarily due to the lack of an appropriate model. Thus, there is a pressing demand for a new microphysiological system (MPS) that preserves LSEC phenotype while capillaries are not embedded -or clamped-within a thick, fixed ECM gel.

In response, we present a vascular engineering technique that successfully guides LSECs towards the spontaneous formation of lumenized microvessels resembling the size and morphology of liver sinusoids. Our approach involves modifying the traditional 2D vasculogenesis assay (see graphical abstract and Fig. 1), which is commonly used as a functional test of endothelial cells at vessel formation ^15,16^. In this assay, cells self-organize into thin, yet stable, interconnected multicellular cords of short lengths when using a reconstituted murine BM coating, such as Matrigel™, GelTrex™ (or other commercial alternative of this sarcoma-derived ECM), as the culture substrate. The composition of these gels—laminin, collagen IV, nidogen, and perlecan—closely imitates the ECM composition of early development^17^ and is pivotal for cell polarization ^18,19^, vessel formation and stability ^15,20^, via key interactions with laminin ^11,21^. Indeed, the underlying mechanisms behind such a cell autonomous organization in a cord network are predominantly mechanical ^22^. It has been recently demonstrated that the interplay between the mechanical plasticity of laminin and the contractility of cells is required for neighboring cells to mechanically sense one another, polarize, and migrate towards each other, ultimately leading to the reorganization of the surrounding ECM to stabilize the structures ^23,24^. Moreover, other culture conditions, including cell density, substrate density and thickness, as well as substrate anisotropy, have been identified in experimental data and models as critical for controlling the final architecture in these vasculogenesis assays ^25^. Through their interactions, these microenvironmental parameters direct the organization of ECs into distinct patterns, including high cellular density spots or aggregates, isolated cords of cells, and cell networks. Particularly, the direction of the organization can be controlled, with migration increasing in the direction of the largest dilatation of the medium in an anisotropic substrate ^26^.

**Figure 1.**
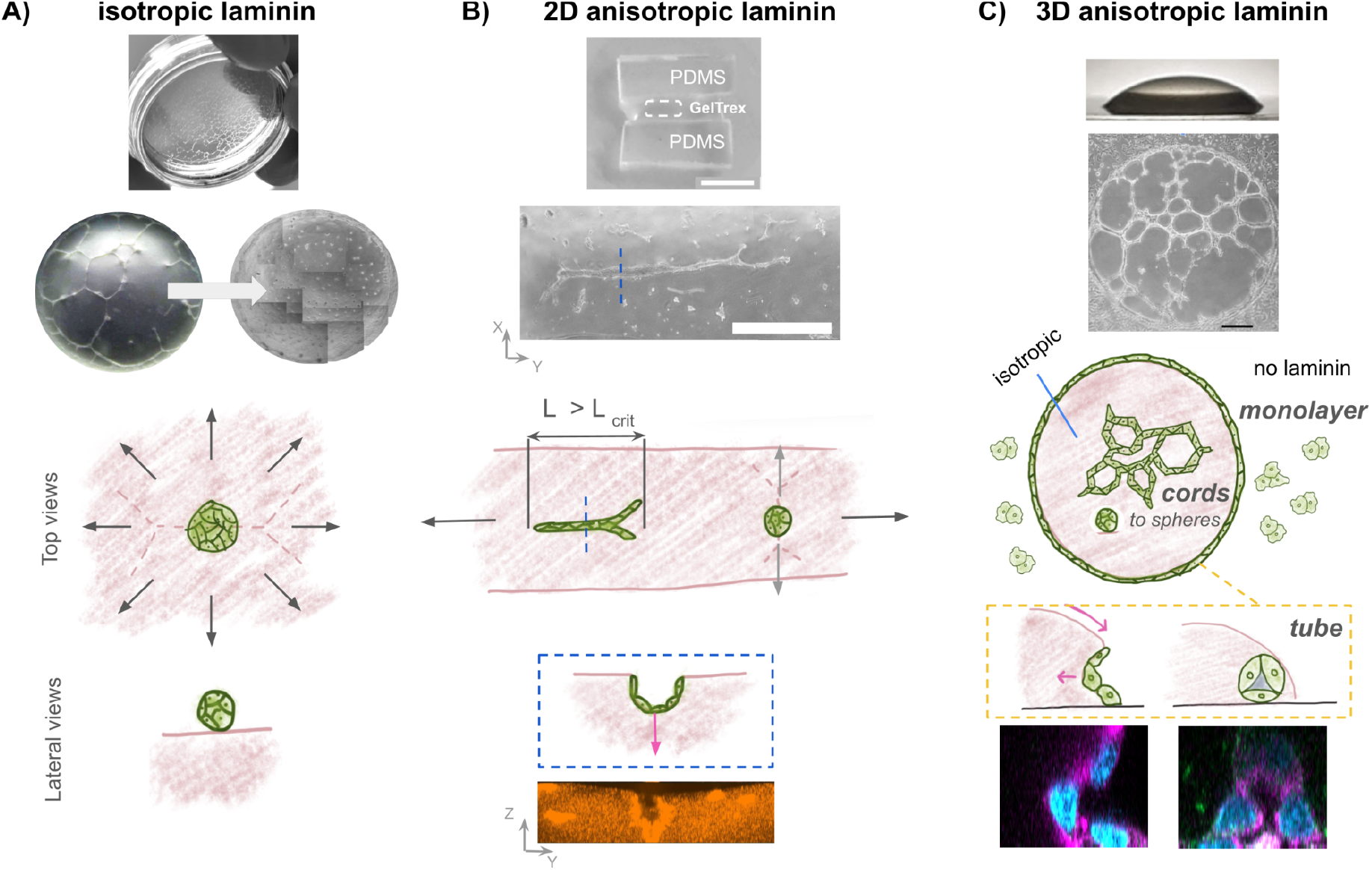
Different geometries of deposited GT guided ECs towards different morphogenetic processes and final architectures. A) The use of flat GelTrex deposits, either in Petri dishes or spread on glass coverslips, guides LSECs to organize into networks with random directions. On this substrate laminin, essential for organization, is isotropically oriented, leading cells to form networks that eventually break down to form aggregates. B) The gelation of a line of GelTrex supported by two PDMS slabs modified the anisotropy of the gel’s aspect ratio. This favored a cord like organization of LSECs in the longitudinal preferential direction. However, when verifying the structure from the lateral view, we observed the cells were deforming the gel in a similar manner as in an invagination but that it has not closed completely. Scale bars: top = 5 mm, center = 1mm. C) We modified topographical guidance using a Marangoni flow-derived patterning, inspired by ^29^. By precise control of the evaporation rate during gelation of the sessile droplet a high protein density can be localized at the peripheral edge. This micropattern successfully confines LSECs in a peripheral train at this edge providing a longer sustained, unidirectional guidance of EC cord organization. Mechanical deformation from the cord along the edge of the dome then triggered a piecewise folding of the soft gel wall, leading to a wave-like blanketing of cells. This process resulted in a complete coating of the cell cord with laminin, facilitating a gradual apico-basal cell polarization and the subsequent cues leading to lumenogenesis. Ultimately, the entire process yielded a lumenized vascular structure around the dome.

We briefly evaluated the organization of LSECs when cultured on GelTrex gelated deposits of different geometries, yielding results consistent with the literature. When using a flat coating of GT, featuring an isotropic deposit of laminin, the cells organized into randomly oriented cord networks. Depending on initial cell density, these cord networks either continued proliferating until forming a monolayer or eventually detached and formed compact spheres, probable outcome of elevated cellular tension in the cords (Figure 1A, Figure S1A). We then engineered thick but flat GT substrates with high length-to-width aspect ratios by using polydimethylsiloxane (PDMS) slabs as scaffolds. We observed that such a geometrical anisotropy guided longer cell alignments in the direction of the gel length, while orthogonal cords or shorter than a certain critical length (Lcrit) of ∼1 mm still resulted in spheres, in accordance with what was modeled in ^26^. When observed in a transverse view, the center of these long-lasting millimeter-ranged cords revealed that the anisotropic gel had been folded by LSECs, resulting in a semi-closed structure that was oriented inwards. However, no evidence of cellular migration outside of the pre-existing cord was observed, nor further maturation of the structure into a vessel (Figure 1B, Figure S1B).

Therefore we focused our study on engineering long and oriented lumenized vessels through the use of gelated sessile droplets of GT as cell culture substrates (Figure 1C). Characterization of the resulting dome showed that the ECM at the middle had an isotropic organization, while having an anisotropic edge given by a thin micropattern of laminin and collagen IV. LSEC organization showcased this regionalization, for they formed networks of cords in the center of the dome and a single, oriented and lumenized structure emerged at the anisotropic edge. Through live imaging we identified the morphogenetic steps that led cells to this structure. First, cells in close proximity to the dome migrated towards the edge, adhering to the ring-like pattern. These cells then followed this trail, undergoing anisotropic migration and proliferation, which gradually organized the sparse, individual cells into a 1-cell ring-like cord (1D) tethered to the thin dome edge. Upon the complete coverage of the edge, proliferation locally changed, resulting in the formation of a multicellular cord (2D) that spread in a planar manner over a distance of approximately two to three cell widths. Mechanical deformation from the cord along the edge of the dome then triggered a piecewise folding of the soft gel wall, leading to a wave-like blanketing of cells. This forcibly surrounded the closed cell tube (3D) with a thin laminin-rich matrix blanket that triggered a lumenogenesis process comparable with the *in vivo* description ^27,28^. A gradual apico-basal cell polarization was demonstrated by the translocation of apical protein Podocalyxin (PDCLX), the maturation of the tight junction was verified through the change of localization of zonula occludens 1 (ZO-1) protein, while mechanical support from the actin cytoskeleton was evidenced by the concentration of F-actin in the newly established apical domain of the structure. Finally, regarding their integration to MPS, we demonstrate that the vessel can be recapitulated when using a straight gel geometry, which could facilitate its cannulation in future work. Notably this simple fabrication method guided an autonomous self-organization of LSECs into a capillary without the need for supporting cells or complex prefabricated scaffolds which is a step towards microphysiological systems of discontinuous, fenestrated capillaries.

## 2. Methods

### 2.1 Glass functionalization

Glass coverslips (18 mm, Ted Pella) were sonicated twice in 96% ethanol, followed by a final round in distilled water. In a fume hood, a 2% 3-(Trimethoxysilyl)propyl methacrylate (TMSPMA, Sigma-Aldrich 440159) (v/v) and 1% Acetic Acid (v/v) solution was prepared in absolute ethanol. Each coverslip batch, with a maximum of 15 pieces, was submerged in 10 ml of the solution in a 10 cm petri dish and agitated for 10 min at medium speed. Excess solution was discarded, and the coverslips in the petri dish were rinsed with 96% ethanol within the hood. Then they were individually removed from the dish, dipped into clean 96% ethanol, and left to air dry. Functionalized coverslips could be stored in a dust-protected container for one week before their use.

### 2.2 Control of gelation of GelTrex™ structures on glass

Functionalized coverslips were placed in a 12-well plate and UV-sterilized for 30 minutes. Various geometries of GelTrex deposits (GT, Gibco, A1413202) were generated. All work performed with GT, prior to gelation, was conducted on an ice bed. In the isotropic laminin condition, a 10 μL drop was placed on each of the functionalized coverslips and then spread evenly over the entire surface. In the 2D anisotropic laminin condition, two polydimethylsiloxane (PDMS Sylgard 184, Dow Corning) slabs were used as a scaffold to deposit a line of 15 μL of GT, with the help of a micropipette. Finally, for the 3D anisotropic laminin condition, a 10 μL drop of GT was added to each coverslip, without spreading it. In all cases, the plate was incubated at 37°C with 95% relative humidity for 30 minutes to promote gelation. Following incubation, 1 ml of medium was added to each well to prevent drying before cell seeding.

### 2.3 Cell culture

Transformed sinusoidal endothelial cells (TSEC, provided by P. Rautou) were maintained in Dulbecco’s Modified Eagle Medium (DMEM, Gibco, 31966021) supplemented with 5% fetal bovine serum (FBS, Gibco, 16000044), 1% Penicillin / Streptomycin (Pen/Strp, Gibco 15140122), 0.4% Amphotericin B (Gibco, 15290026), 1x Endothelial Cell Growth Supplement (EGCS, CliniSciences, 1052-SC). TMNK 1 cells (provided by M. Macías-Silva) were maintained in DMEM, High Glucose, GlutaMAX™ (Gibco, 10569010) supplemented with 10% fetal bovine serum (FBS, Gibco, 16000044), 1x Antibiotic - Antimitotic (AntiAnti, Gibco, 15240112). Following trypsinization, cells were counted using a CASY cell counter (OMNI Life Sciences) and cultured at a density of 196 cells/mm^2^, ensuring even cell distribution.

### 2.4 Scanning Electron Microscopy

Samples were fixed with 2% glutaraldehyde (Sigma-Aldrich, 340855) in 0.1M cacodylate buffer pH 7.4 for 2 hours at RT. Then samples underwent five 10-min washes in gentle agitation with 0.1 M cacodylate buffer. Post fixation was performed by incubating samples on ice for 20 min in 1% Osmium tetroxide in 0.1M cacodylate buffer. Samples were washed for 5 min in gentle agitation with distilled water, five times. Dehydration was achieved by subjecting samples to 10-min incubations with increasing concentrations of ethanol 50%, 70%, 95%, 100%, absolute ethanol. Final drying was performed through evaporation of hexamethyldisilazane (HMDS), first with a 10 min incubation of HDMS 50% in absolute ethanol, followed by two incubations of HMDS 100%. The dried samples were mounted onto sample holders and coated with platinum before being imaged with a Cryo-SEM GeminizeM500 using immersion lens and SE2 detectors. We used a High resolution GeminiSEM 500 (Zeiss) for the scanning electron microscopy (SEM).

### 2.5 Filling factor determination

SEM images from the center, near edge and edge of the dome were obtained as previously described. Pore size and occupied area were determined using a threshold to select the void spaces. The filling factor was computed as the ratio between the area occupied by the matrix and the total field area. Results are presented as percentages.

### 2.6 Nanoindentation of GelTrex domes

The Piuma system from Optics11 was used to perform nanoindentation measurements of GelTrex gelified deposits immersed in complete medium. Load-displacement curves from the bending of the cantilever were obtained over the course of several dome radii. Estimation of the material’s Young’s modulus was performed using a Hertz model fitting.

### 2.7 Phase contrast live imaging

35 mm glass-bottomed petri dishes (World Precision Instruments, FD35-100) were functionalized with TMSPMA as previously described. Subsequently, a 10 μL GT droplet was placed in each dish, gelated and cultured as described in 2.2. Phase contrast microscopy images were acquired every 10 min for 5 days using a BioStation IM-Q (Nikon). Particular attention was placed on the edge of the dome, leaving the central part of the dome out of the study. From the generated videos (see supplementary material), displacement and angle of displacement were obtained through Tracker Video Analysis and Modelling Tool ^30^.

### 2.8 Immunofluorescence

Samples were fixed for 15 min at 37 °C with 0.4% Paraformaldehyde (PFA. Sigma-Aldrich, P6148) in Phosphate-buffered saline (PBS, Sigma-Aldrich, P3813), permeabilized with 0.1% Triton X-100 (Sigma-Aldrich, X100) in PBS for 10 min at RT, in agitation; and blocked with 10% horse serum (HS, Sigma-Aldrich, H1270) for 1 hour. After blocking, primary antibodies corresponding to Laminin 111 (Novus Biologicals, NB300-144SS) (1:1000 dilution), Collagen IV (Novus Biologicals, NB120-6586SS and NBP1-26549) (1:500 dilution), Podocalyxin (R&D systems, AF1556) (1:250 dilution), and ZO-1 (Invitrogen, 10342463)(1:50 dilution) were diluted in blocking buffer and incubated at 4°C overnight. Then samples were incubated for 2 hours with second antibodies: goat anti-rabbit (Alexa Fluor 633, 1:500), donkey anti-goat (Alexa Fluor 594, 1:400). For F-actin staining, samples were incubated for 45 min with Alexa Fluor™ 488 Phalloidin (Invitrogen, A12379). Nuclear staining was performed through incubation of 4′,6-Diamidino-2-phenylindole dihydrochloride (DAPI, Sigma-Aldrich, D8417) diluted 1:250 in PBS. Samples were imaged with a Zeiss LSM 980 Inverted confocal microscope using an oil-immersed objective (63x) with a 0.5 µm step for Z-stack acquisition.

### 2.9 Fluorescence quantification

Image analysis was performed using Fiji image analysis software ^31^. Pixel intensity was obtained through profile plotting either of straight selections or oval selections ^32^. For each experiment, intensity was normalized to maximum intensity value.

For the quantification of ZO-1 condensate, using images from stage 3, a threshold was determined to select only the clusters. Using this threshold to generate binary images in both stages evaluated, the ‘*analyze particles*’ command was used to obtain the number of condensates.

### 2.10 Fluorescence live imaging

For live fluorescence microscopy, FluoSpheres™ Carboxylate-Modified Microspheres (Invitrogen, F8786) (dilution 1:1000 in DMEM) were added to the GelTrex prior gelation (final GT concentration 95%). Additionally, cells were incubated with CellMask™ Green Actin Tracking Stain (Invitrogen, A57243) (dilution 1:1000 in starving medium) for 30 min at 37°C. Nuclei were stained with Hoechst (Invitrogen, 33342) (diluted to 1 ug/mL) for 10 min at RT. Image acquisition was performed using a Zeiss LSM 980 Inverted confocal microscope. Pictures were taken every 30 min.

### 2.11 Displacement map generation

Z projection stacks obtained from live imaging were first aligned to correct drift, using the “Linear stack alignment with SIFT” plugin from Fiji. We divided the complete stack into sets of 2 images that showed the evolution of the cell culture (Picture_i_, Picture_i+1_). Then, for each stack, we ran the iterative analysis from the Particle Image Velocimetry (PIV) plugin ^33^. As suggested by the developers, we established the interrogation window to be ¼ of the picture size and the search window to be ‘interrogation window’ size plus twice the maximum displacement. And for each iteration search window size was set to half of the previous value. Displacement fields for each time step were obtained through the plot instruction of the plugin. To obtain the complete trajectory of each bead, the complete stack, obtained in 2.10, was analyzed through the TrackMate plugin ^34^.

### 2.12 Statistical analysis

If not stated otherwise, data are presented as mean ± SD from at least two independent experiments. Graphs were generated through the Matplotlib graphic package ^35^. Statistical analysis to assess differences among experimental groups was performed using Python’s SciPy sub-package ^36^. We first assessed the assumption of normality and homogeneity of variance for each dataset. Based on the results, we applied an appropriate statistical test to determine significant differences, which are indicated under the corresponding figure. A significance level of P<0.05 was considered significant and notation is as follows: P<0.05 “*”, P<0.01 “**”, P<0.001 “***”, P<0.0001 “****”.

## 3. Results

### 3.1 Gelation of a sessile droplet of GelTrex densifies the deposition of Laminin and Collagen IV in a ring-like pattern at the edge

The first step of our process for non-encapsulated microvessels consists in guiding the cultured endothelial cells to form long, thin cords that serve as a foundation for alignment for the formation of a vessel with lumen. As mentioned in the introduction, we engineered a laminin-collagen IV micropattern using GelTrex (GT) to guide cell adhesion, proliferation and migration in one preferential direction. Indeed, laminin is key in driving EC cord formation, on the one hand, and in controlling EC apicobasal polarization responsible for lumenogenesis, on the other hand. This thin pattern was formed at the edge of a hydrogel dome geometry, as described in the methods section (Figure 2A). Similarly to what was reported with domes of collagen I^29^, the ECM was distributed unevenly along the radial direction in our domes, identifying three zones by SEM: middle, near edge and edge (Figure 2B). We quantified the matrix surface filling factor to assess ECM density (Figure 2C) and found the highest value at the edge (98.13% ± 1.02%), and a significantly lower number in the central middle region (68.17% ± 5.58%) (Figure 2D). This gradually increasing density towards to border of the dome correlated with smaller diameters of the identified pores in the matrix; much smaller at the edge (51.46 nm ± 8.33 nm) than those in the middle of the dome (357.80 nm ± 136.78 nm). These results indicate that the Marangoni effect engineered by Nelson’s group with collagen I to obtain a gradient of ECM density across the radial axis of a sessile hydrogel droplet is not restricted to fibrillar materials and can also be obtained in the gelation of a laminin-rich GT dome.

**Figure 2.**
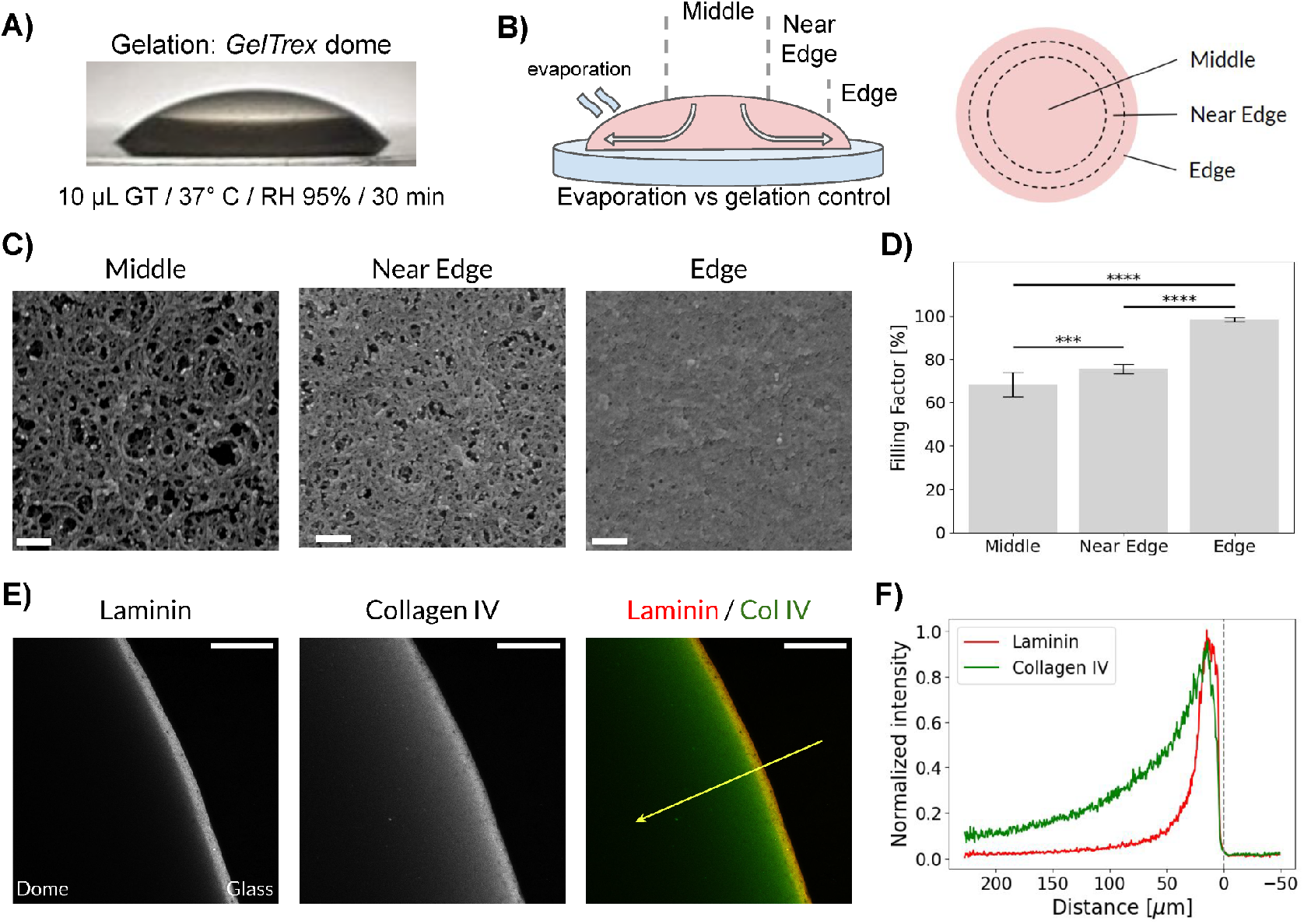
Gelation of a sessile droplet of GelTrex defines a dome with density gradient ending in a ring pattern of laminin at the edge. A) Transversal-view brightfield image of a gelated GelTrex dome. B) Schematic of the evaporation during gelation in Marangoni flow, creating three specific concentric zones: middle, near edge and edge. C) Scanning electron microscopy micrographs of the ECM from the different zones after gelation, increasing its density towards the edge. Scale bars = 100 nm. D) Comparison of computed ECM filling factor in the three zones. Data were analyzed using ANOVA tests and a post-hoc test (Tukey-Kramer test) to determine significant differences between groups. (n=3 per group, “^***^” P<0.001, “^****^” P<0.0001). E) Confocal Z projections of dome edge stained for laminin and collagen IV showing a thin ring-like pattern. Scale bars = 100 µm. F) Intensity profiles of transversal lines pointing inside the dome (yellow arrow) traced for laminin and collagen IV. Average profiles for 4 samples for each stainings; distance 0 µm is the dome edge.

We also performed mechanical tests using nanoindentation that aligned with the observed increase of ECM density, revealing a radial increase in the Young’s modulus towards the periphery. Specifically, a value of (245.5 Pa ± 40.3 Pa) was found at the center of the domes and drastically and locally increased to (439.5 Pa ± 156.2 Pa) at the very edge. Both of these measurements are consistent with the reported elastic modulus from other reconstituted BM gels, for example the (450 Pa ± 230 Pa) for Matrigel ^37^. Finally, we examined the distribution of laminin and collagen IV after gelation by using immunostaining (Figure 2E). Both proteins exhibited a distinct high concentration in a ring-like pattern at the edge of the dome, a characteristic result of Marangoni flows ^38^. Their profiles were analyzed transversally and inwards through the dome (yellow arrow in figure 2E) and the full width at half maximum (FWHM) was measured for both signals. A thinner (16.78 μm ± 3.89 μm) laminin signal was revealed compared to that of collagen IV (44.75 μm ± 4.54 μm), as shown in Figure 2F, where the edge of the dome is defined as the origin. Importantly, the dense laminin trace can be considered a micropattern, for guidance, given its order of magnitude relative to a single cell size. Previous studies have shown that such narrow patterns could guide EC spreading, orientation and migration ^39^. This is particularly relevant to our efforts in confining cell growth and migration on laminin to align multicellular cords along one single, anisotropic path. Remarkably, and as noted in ^29^, the correct sample preparation and environmental conditions during dome gelation conditions detailed in the Materials and Methods sections are of the greatest importance to obtain this laminin pattern. We present some failed conditions in the supplementary material (collagen instead of GT, spreaded dome with low contact angle, a different culture medium, a stiffer gel, a greater cell density), to help the readers reproduce our experiments (supplementary text, Figure S2).

To avoid the encapsulation of vessels constructed by the hepatic endothelial cells and obtain an architecture that resemble more closely that of a liver sinusoid with only a thin BM, we demonstrated that the edge of the dome structure could guide LSECs to self-organize into a surface-bound cord aligned in a preferred direction (as opposed to the random cord pattern), then fold the gel and close a tube with a lumen. The 3D geometry offered by the dome substrate with a dense laminin pattern of laminin and collagen IV at the dome edge contrasts greatly with the uniform, flat ECM layer employed in traditional vasculogenesis assays (Figure S3A).

### 3.2 Liver sinusoidal endothelial cells form tubular structure following the edge of a GelTrex dome

Given this 3-dimensional organization, and following our observations where anisotropy can force cells to fold their substrate, immortalized liver sinusoidal endothelial cells (TSECs) ^40^ were then cultured on the dome samples and their organization was followed over time. Three distinct types of cell structure emerged, depending on the localization in the sample, as depicted in Figure 3A. Outside the dome, TSECs that adhered to the glass rapidly formed a monolayer surrounding the dome. In the middle region and near-edge region of the dome, cells were organized into a network of cords resembling the structures formed in traditional self-organization assays performed on flat Matrigel or GelTrex ^16,24^. They first organized into isolated cord fragments with an average length of 238.4 μm ± 75.6 μm and an average width of 10.5 μm ± 1.6 μm. Then, the number of cords increased almost 5 times to obtain an interconnected network. Without increasing their average length (240.1 μm ± 87.4 μm), cords became thicker in the middle region, with an average width of 22.1 μm ± 6.5 μm. At 96 hours, there was a remodeling of the network shown by the decrease in the number of cords by half, the decrease in the number of interstitial spaces, and the increase in average cord length (372.6 μm ± 84.7 μm). Finally, a tubular structure of interest was formed by the cells that attached to the edge of the dome (zoomed region of Figure 3A). As detailed in later sections, these cells populated the edge came from the neighboring glass and oriented along the laminin micropattern (Figure 3B and Supplementary Figure 4A) to gradually form a piecewise train of cells (Supplementary Figure 4B). This specific 1-cell wide cord type had an average width of 11 μm ± 2.3 μm that was sustained for longer than the other cords. The different peripheral pieces of cords presented varied lengths, ranging from 170 μm to 1900 μm to cover 60% of the circumference of the dome after 24 hours of culture. After 48 hours, they had merged into a single cord covering the entire periphery of the dome, connected with the network formed at the center of the dome in a few contact points, Figure 3A. After 72h, it was clear that long portions of the edge looked like 3D tubes, while the other structures on the dome remained thick but flat, and immunostaining for VE-cadherin confirmed the presence of a lumen (Figure 3C). Furthermore, at 96 hours, an ECM front, that was gradually moving away from the original edge of the dome (indicated with a yellow arrow in Figure 3A) becomes more visible as it connects with the external monolayer. It was found that it is a small portion of the GT dome displaced by migrating cells and its progression was quantified during the whole process of tube formation (Figure 3D). We detail in a later section that it plays an important role in lumen formation (Figure 8B). Interestingly, if the conditions of initial droplet formation and gelation were not carefully respected and a monolayer was allowed to invade the edge before an early cord was formed, no tube could be formed (Supplementary Figures S2B and S2E).

**Figure 3.**
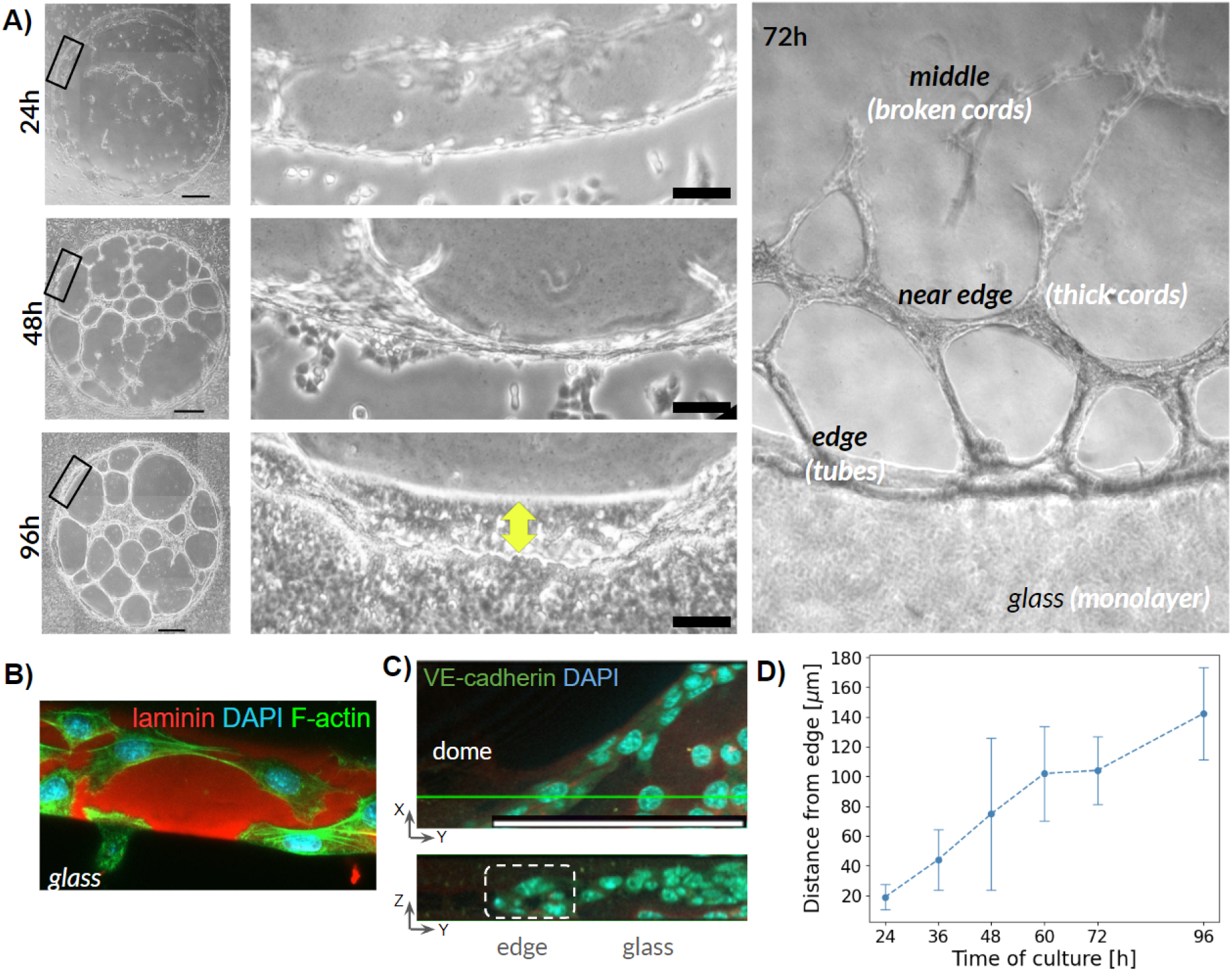
Endothelial cells self organize in a lumenized structure following the ring pattern at the edge of a GelTrex dome. A) Top-view brightfield images of TSECs cultured on GelTrex domes at 24h, 48h and 96h. Left side pictures allow to visualize the whole dome with networks forming in the center, as conventional vasculogenesis assays, scale bar = 500 µm. Black rectangles indicate the zoomed region shown besides, focusing on the edge and showing the progression of the organization, scale bar = 100 µm. B) Confocal image of organizing cells. Cells on the gel are migrating towards each other in order to form the cords while at the edge (bottom left) a cell can be seen adhering to the dome. C) Confocal micrograph of the structure formed at the edge of a dome, scale bar = 100 µm. Below, transversal cross-section showing a tubular structure with lumen (dashed square). D) Measured distance from the edge of a front formed during the cell organization, showing an increase in width, n = 3.

We characterized all these different cell-made architectures after 96h in culture through SEM. As expected, it revealed distinct multicellular morphologies and cellular ultrastructures adopted by the TSECs in these three regions. A cobblestone morphology, characteristic of confluent monolayers, was visible outside the dome (Figure 4A). These cells presented a high density of microvilli, structures normally associated with pathological LSECs ^41^, confirming that a monolayer is not representative of an endothelial vessel phenotype. The cords in the middle and near-edge regions of the domes appeared to be multicellular structures composed of slightly curved cells that had no microvilli, like piled up tiles and in the process of forming tubes (Figure 4B). They exhibited zipped, loose cell-cell junctions, not mature for a vessel, with clear openings between neighboring cells. Moreover, submicron-sized *fenestrae*, indicative of a healthy LSEC phenotype (and typically absent from LSECs in culture after 1-2 days), were also observed on these cells (zoomed in Figure 4B). However, the cords appeared to be only sitting on top of the ECM, with no real interaction with the underlying ECM, contrary to what was described by others during EC vasculogenesis on Matrigel ^24^, where cells organized the surrounding matrix to stabilize the cords they constructed. In contrast, the peripheral structure was organized into a more elongated and thinner tubular structure over the whole dome edge (Figure 4C). The formation of this structure is presented in Figure S4. Compared with the central cords, this structure appeared to be more connected with the ECM and covered by a thin BM after 96h, as no cell membrane or cell-cell junction were visible under it. As suggested by other studies ^24^, this BM was probably secreted by the cells: interestingly, we observed that over time, this coating thickened and aligned along the tube, demonstrating vessel maturation (see Figure 9E). Importantly, this regionalized organization was also obtained with an immortalized human LSEC cell line (TMNK-1 cells) ^42^, following the same process, demonstrating that this process is not a cell line-specific feature (Figure S5). These results show that the engineered sessile droplet of GT, in a soft dome-shape substrate, instructs LSECs to self-organize differently depending on ECM density and organization. Thanks to LSEC high contractility ^43^, cord networks also formed in areas with randomly organized ECM at the center of the dome, similar to conventional vasculogenesis assays. Regions along the edge exhibited a more stable, vessel-like morphology with a preferential orientation.

**Figure 4.**
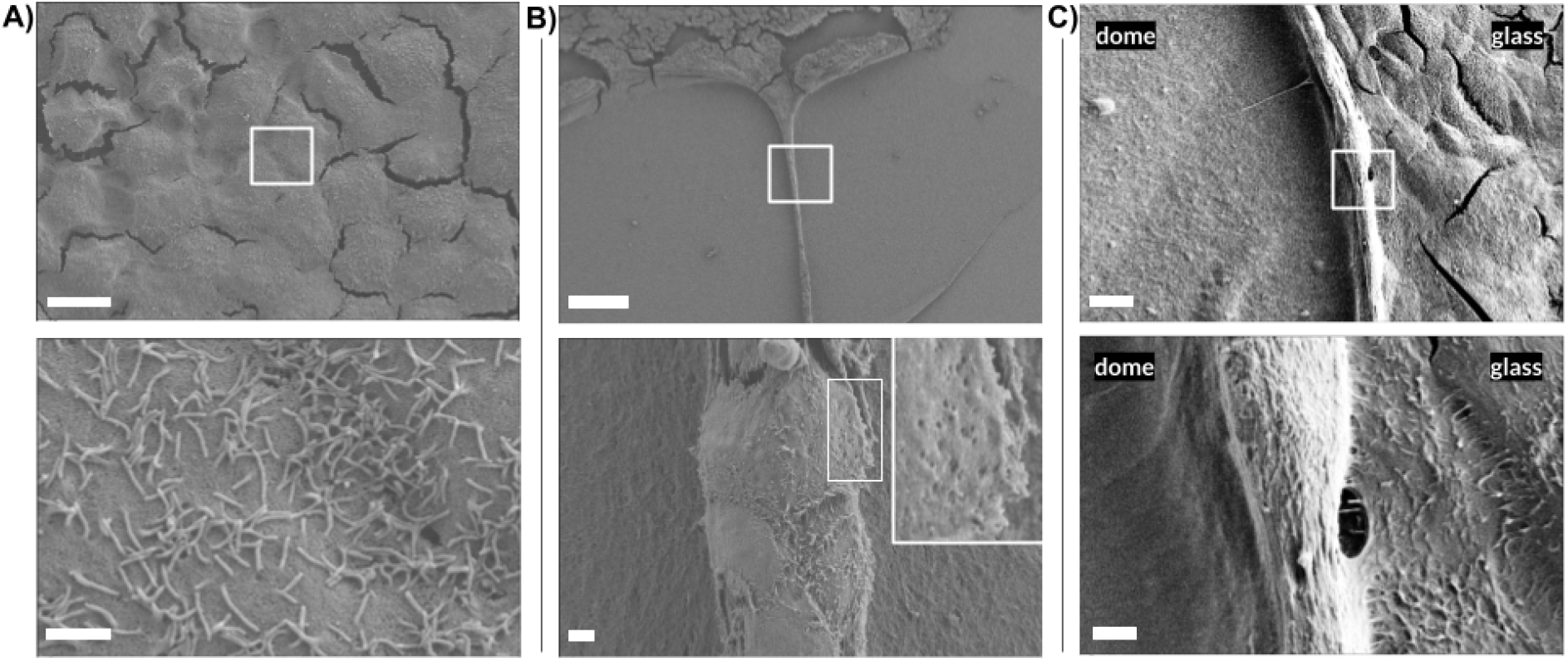
Culture on GelTrex domes guide endothelial cells into three different morphogenetic multicellular shapes depending on their localization. Representative micrographs from scanning electron microscopy of TSECs cultured for 96 hours on a GelTrex dome. A) Cells attached to the glass proliferated and formed a monolayer characterized by the cobblestone morphology, scale bar = 10 µm. Zoomed region shows the presence of cilia structure on the cells, scale bar = 1 µm. B) On the dome, cells are organized in multicellular cords that are connected to the edge, scale bar = 100 µm. Zoomed region shows loose cell-cell junctions and large openings, scale bar = 2 µm. C) Elongated cells at the edge (dome left, glass right) organize in a cord, scale bar = 10 µm. Zoomed region shows the cord covered by a thin ECM-like coating, scale bar = 2 µm.

### 3.3 Progressive morphogenetic steps at the edge of the GelTrex dome enable a lumenized microvessel formation

Further experiments were conducted to elucidate the precise morphogenetic steps in the dynamic behavior used by TSECs to construct the tube. This section details the four consecutive stages that were identified as crucial to guarantee the generation of the tubular structure with a preferred tangential direction at the edge of the dome. We show that the initial steps depend on the specific geometry of the GT dome and that the lumen formation was entirely constructed by the cells, in response to 3D cues of the surrounding laminin polarization.

#### Stage 1: formation of a long, 1-cell thin train along the dome edge

Since the earliest moments of culture, cell dynamics was controlled by preferential migration/proliferation patterns. First, once adhered to the surrounding glass, cells in a close proximity began migrating towards the edge of the dome (Figure 5A, Movie S1). We tracked cells progression during the first day of culture in the vicinity of the dome and measured their gradual approach (Figure 5B, Figure 5C). A threshold distance of approximately 55 µm, equivalent to 4 cell sizes, was observed: cells located below this distance migrated toward the dome at an average (and relatively constant) speed of 2 µm/h, while cells further away did not get closer and even moved away. This approach migration speed seems low compared to reported values *in vitro*. We recall that no ECM was deposited on glass prior to culture. It is, however, of the order of magnitude of velocities found for HUVECs in similar low density conditions ^44^. The direction of migration during this approach phase was evaluated by measuring the angle between the tangent to the dome edge and the direction of migration. Before attachment, no specific preferred angle was found: Figure 5D shows a broad distribution of approaching angles per cell, ranging from 6° and 162°. However, after attachment to the dome edge, cells aligned along the edge (as depicted in Figure 3B) and the direction of migration remained anisotropic, permanently along the edge, in a 1-cell wide train. This is shown in Figure 5E, presenting cell orientation along an angle of either 180° or 0° depending on the clockwise or anticlockwise direction of the migration immediately after contact. Moreover, the anisotropic direction of migration was persistent and maintained by a preferred direction of division which followed the mother cell’s previous orientation and helped construct the piecewise cell train, as depicted in the representative examples of Figure 5A for both approach (white arrows) and edge adhesion (yellow) phases. Importantly, the structure self-organizing at that stage on the gel remain isolated and disconnected from the monolayer forming outside (Figure 8C).

**Figure 5.**
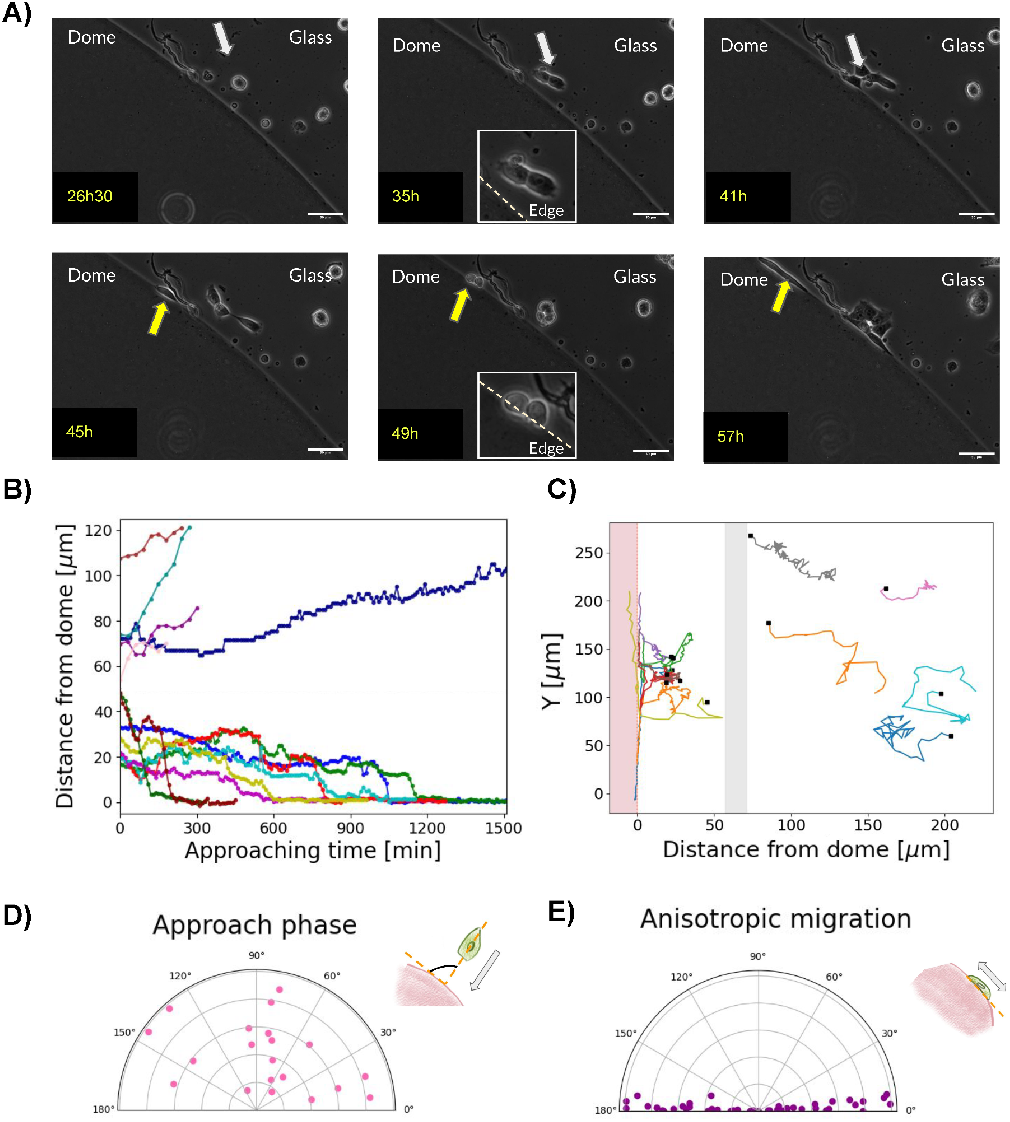
First stage of endothelial organization is characterized by migration towards the dome and preferential adhesion and proliferation along the edge. A) Micrographs from bright field microscopy (frames from MovieS1) showing the progression of organization at the edge. First, some cells migrate towards the dome (white arrow, top line) and proliferate with no particular direction (top). After reaching the edge, they modify their direction of migration and proliferation anisotropically along the edge (yellow arrows, bottom line). Scale bar = 50 µm B) Quantification of the approach phase of individual cells. Minute 0 corresponds to the start of cell displacement n = 13. C) Representative trajectories of individual cells on the substrate. Starting point is shown by a black square for each cell. Cells closer to a ∼50 µm distance threshold (gray zone) migrated towards the dome. D) and E) Angle between the direction of cell migration and the tangent to the edge confirmed the observations in A). n = 8.

#### Stage 2: formation of a multicellular cord around the dome

Once adherent to the ECM, TSECs migrated and proliferated exclusively along the dome edge, in the tangential direction, to gradually cover the entire edge (Figure 6A, Movie S2). During this second day, cell speed of migration increased to an average of 7 µm/h; they also stretched greatly while migrating and before dividing. Between two to three days after seeding, the full dome was bound by the cell train before a gradual change was found in the structural organization. Some cells were indeed found at a different height, corresponding to the start of cell proliferation outside the circumferential axis of the dome edge along which they were previously confined. Then, the edge clearly deformed in a piecewise fashion, suggesting a cell-driven process.

**Figure 6.**
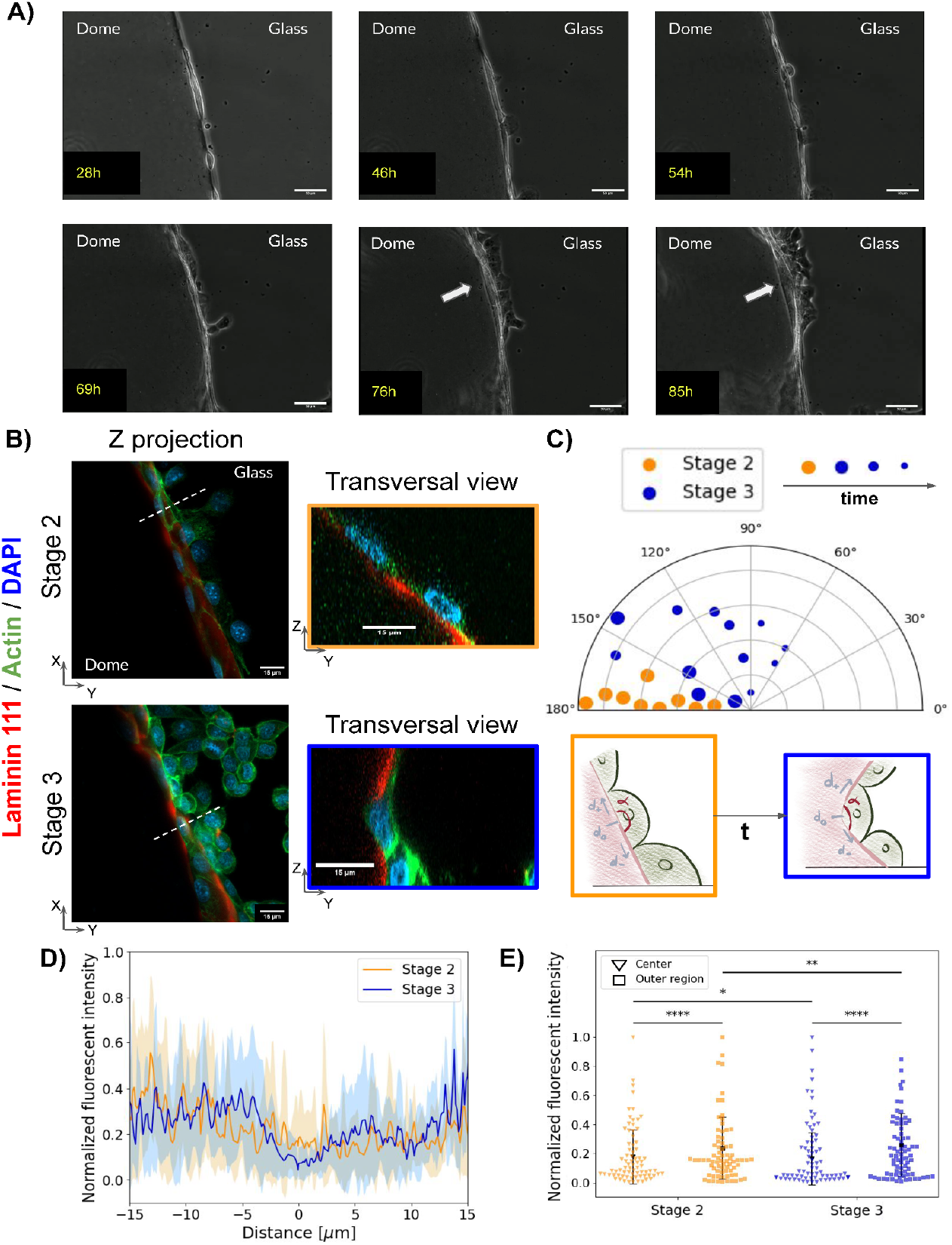
Progression of the thin cell line into a multicellular cord structure and ECM edge folding. A) Micrographs from bright field microscopy (from Movie S2 and Movie S3) showing the cord formation at the edge. Top line: cells continued to migrate and proliferate around the edge until covering the whole circumference with a thin cord. Bottom line: Cord thickened, with the appearance of a white front, moving away from the dome (white arrow). Scale bar = 50 µm. B) Two stages were identified with Z projections of both stage 2 and 3. Scale bar = 15 µm. Transversal cross-sections show that the cells folded the dome at a hinge cell, where laminin seemed locally depleted. Scale bar = 15 µm. C) The folding angle represented in the schematics, between the upper cell (in the *d*_*+*_ direction) and lower cell (in the *d*_*-*_ direction), centered at the hinge cell in *d*_*0*_, was measured for stage 2 (orange) and 3 (blue dots with decreasing size for longer times). n= 3 and 5 per group. D) Laminin intensity profiles were measured at the edge for stage 2 and 3 and show an apparent laminin depletion in the central region (around *d*_*0*_ *=* 0 µm) while this effect is reversed at 5 µm in negative (*d*_*-*_) and positive (*d*_*+*_) directions and no intensity difference was found beyond. n= 3 and 5 per group. E) Comparison of normalized intensity at the center and outer region (lower cell d-). Data were analyzed using Kruskal-Wallis tests and a non-parametric *post-hoc* test (Mann-Whitney U test) to determine significant differences between groups. (n= 3 and 5 per group. “^*^” P<0.05, “^**^” P<0.01, “^****^” P<0.0001).

#### Stage 3: formation of an ECM folding pattern

To examine this process more closely, we stained cells with phalloidin and performed an immunofluorescence against laminin 111, the principal component of GelTrex. Thanks to confocal microscopy, we observed transversal views of samples in this new stage and compared with stage 2, both presented in Figure 6B. We confirmed that the architecture transitioned from the 1-cell train structure (1D) at the edge towards a thicker, 2D multicellular cord that followed the same orientation around the entire dome. We found that the cord consisted of a 3-cell thickness, with cells being more spreaded out at the bottom of the gel and more elongated above, as already seen in brightfield microscopy (Figure 6A, white arrow). This thickening was also associated with a change in the direction of proliferation and resulted in a new train of cells forming further up the edge. This process was observed all along the dome edge (Figure 6A, bottom line) and eventually led to localized and piecewise ECM displacements, which are characteristic of this stage 3. Immunofluorescence imaging showed that this resulted from the local folding of the gel, driven by a local indentation of the vertical edge of the dome. This folding consistently occurred beneath a central “hinge cell” (Figure 6B, stage 3), and we quantified the progression of the folding angle during the indentation (Figure 6C). This angle gradually decreased from a flat substrate (180° at stage 2) to an impressive 60° folding, after 24 hours from the start of stage 3. Importantly, this angle contributed to both cell and dome folding along relatively long sections (that eventually connected to circle the full dome), similarly to an ocean surf wave progressing outwards to the shore. Following this analogy, at smaller angles of the later folding instants, the plunging break rapidly converted into a collapse break (wrapping the cells with ECM), making it challenging to measure the progression of folding accurately, as shown later in Figure 8D.

In biological tissues, folding processes driven by cells have been mainly attributed to the constriction of an apical actomyosin network. However, in our observations, we could not find any indication of such an active cellular process driving the folding. We found a local reduction of the ECM density was consistently shown at the basal side of the hinge cell, precisely where the angles were measured (Figure 6D-E), as a decrease in the laminin signal below the hinge cell, when compared to its neighboring cells. This contrast was even more significant as the folding progressed: starting with a 25% difference and increasing up to 36% at the latest observable instant of stage 3. This observation of a localized ECM depletion at the basal side of the hinge cell is in line with a previous report of tissue-wide folding of epithelial cells, responsible for the developmental morphogenetic process underlying the formation of the Drosophila wing ^45^.

In our study, the folding process occurred around the entire dome and it happened in a piecewise fashion, with all the pieces progressing and connecting to each other after a few hours. However, it was independent of the dome size, from small to large domes tested in the 5 - 80µL volume range (not shown). In order to elucidate if the dynamics behind this folding were due to tissue deformations under cell stress, we performed live-imaging analysis of this 2D-to-3D process on GelTrex domes incorporated with fluorescent beads (Figure 7 shows a representative experiment and analysis). This allowed us to track the displacement of the beads every 30 min, starting after the cord had formed at the edge of the dome and beyond ECM folding. Figure 7A shows the temporal evolution of the displacement of the beads and its analysis shows that the observed folding was a passive deformation following a lateral strain pulling a thin portion of the gel on the outside. Two distinct regions of interest stood out during the experiments, as depicted by the groups of displacement vectors. First, localized lateral displacements appeared in locations where the cord had thickened due to local outwards proliferation of cells (purple boxes). These regions appeared almost periodically (every 300 to 500 µm, delimited by white dashed lines) and were responsible for the piecewise nature of the process. Between them, a third, central region (green box) was defined, where bead displacements always occurred posterior to the lateral ones, with no apparent proliferation, suggesting a collective movement of “follower” cells driven by the two boundaries created by the “leader” cells of the other two regions. Indeed, the lateral strains that appeared first on the two ends showed a lateral and tangential direction, almost symmetrical with respect to the central region where folding then appeared. The bead displacement of the central region occurred 2 hours after the lateral stretches and presented an overall inwards direction, resembling a retraction, with a larger amplitude than the lateral ones (Figure 7B). This central inwards folding event actually corresponded to an ECM blanket covering the cells and curling back inside (Figure 7D), explained by the bead displacement at the central region. A persistence image was finally generated and confirmed this global displacement, as presented in Figure 7E. It provides a comprehensive overview of the overall process. The supplementary material offers a simplified illustration of the proposed folding process, which serves as a visual aid for understanding the underlying principles. Stage 3 can thus be characterized as a cell-provoked, tension-driven folding where the laminin-rich ECM slowly blanket-rolls the dome edge where cells are aligned as a multicellular cord, first by covering it and then curling back inwards.

**Figure 7.**
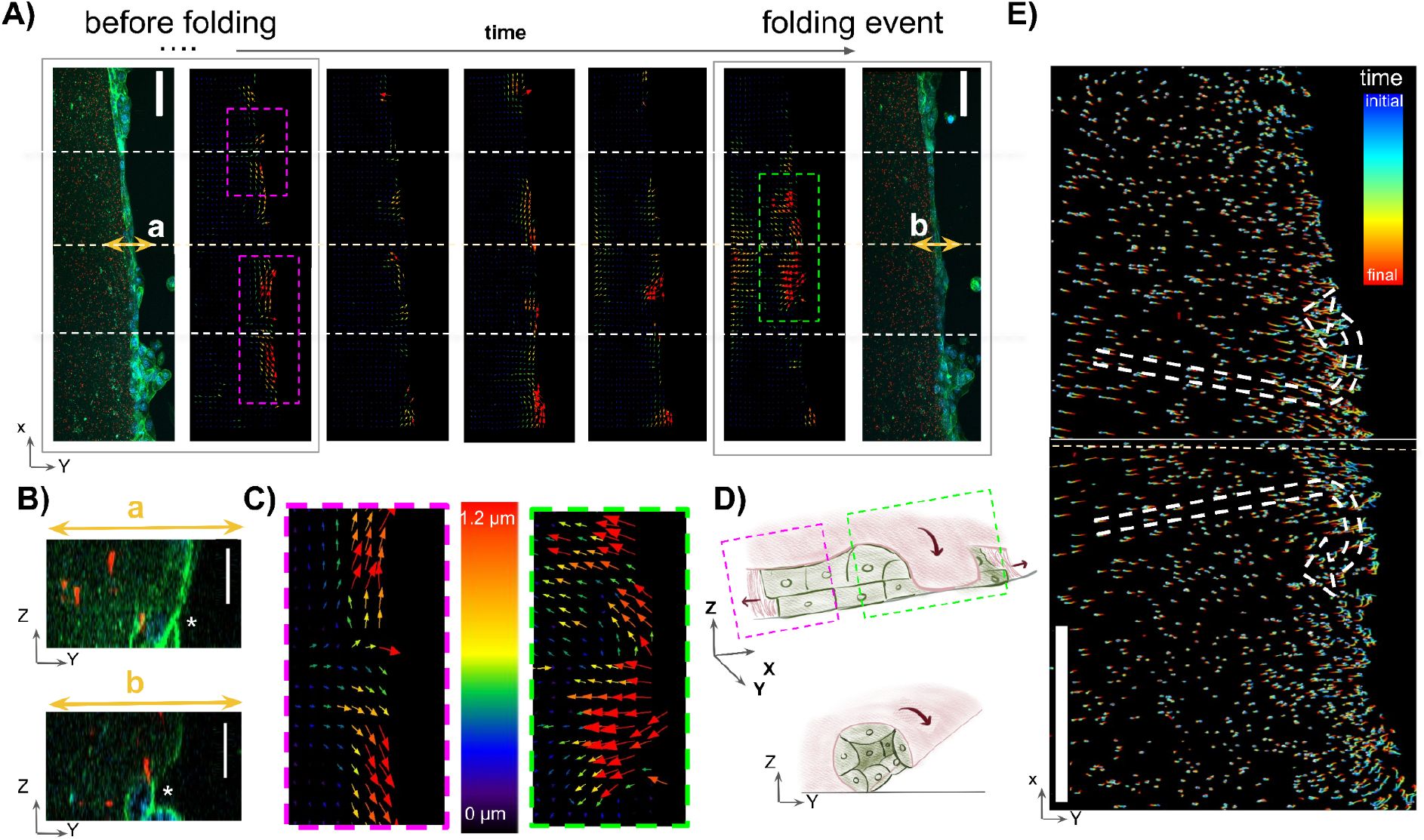
The deformation of the gel in opposite directions along the edge axis initiates folding in a wave-like manner. A) Characteristic temporal evolution of the folding. Confocal images show the edge of a dome with embedded fluorescent beads and cord organization, together with displacement maps. Scale bar = 50 µm. In regions marked as ‘a’ and ‘b’, transverse sections are displayed in B), illustrating the progression. Scale bar = 15 µm. Photographs were taken every 30 minutes. C) A magnified view is provided for characteristic regions displaying lateral bead displacement before folding (highlighted in purple) and inwards bead displacement at the folding event (highlighted in green). Color scale indicates the displacement length. D) Schematic representation of the folding. Cells first deform the gel laterally in opposite directions, leading to an inwards curling front responsible for folding, similar to a breaking wave. The cell cord ends up located inside the wrapping ECM. E) The comprehensive trajectory of each individual bead throughout the experiment is presented. Maximum displacement is observed in the central region of the field, coinciding with the folding event. Color map: blue is the initial starting point of measurement and red the final location. Opposite global directions are clearly visible between the edge and inner regions. Scale bar: 100 µm.

#### Stage 4: ECM enclosure of the multicellular cord and vessel maturation

The final morphogenetic step of our microvessel engineering process corresponds to the completion of lumen formation via ECM folding and surrounding of the cell cord as a blanket-roll shape (Figure 8 and Figure S6). This event is simultaneous with a continued ECM displacement away from the dome edge, sliding out like a wave landing on the shore (Figure 8A). This displacement is correlated with the structure identified in figure 3B. A vessel-like structure established at the edge around the entire dome and featured a capillary-sized architecture with only 3 or 4 cells facing each other, within an ECM roll. The organization of this structure was evident, as indicated by the staining of tight junction protein Zonula occludens-1 (ZO-1), podocalyxin, and actin (Figure 8, Figure S7).

**Figure 8.**
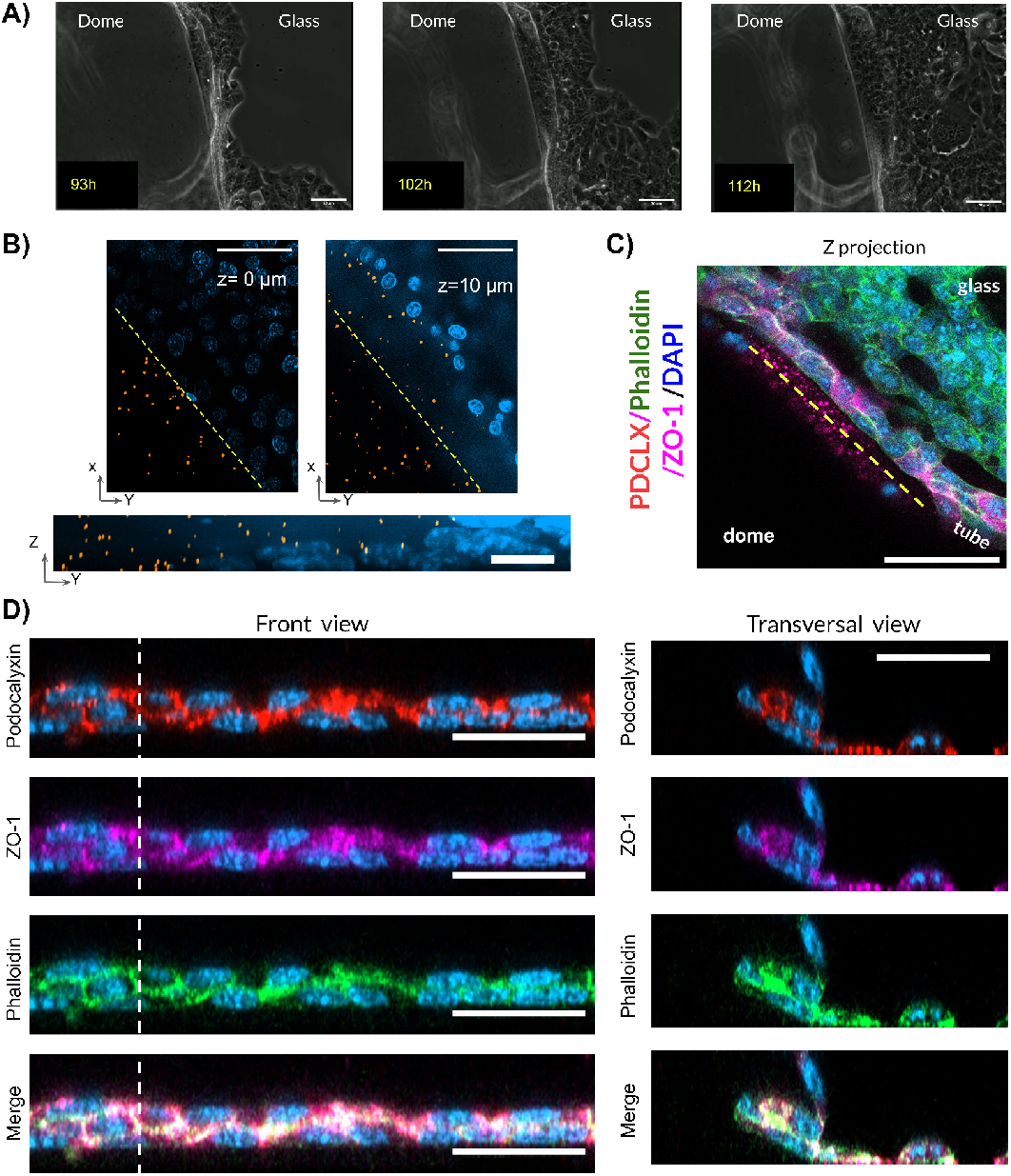
Final stage of organization includes closing of the fold and covering the cells closer to the dome, both cord and monolayer, with gel. A) Micrographs from bright field microscopy (from Movie S4). The cord observed in stage 3 continues to move away from the original edge. Scale bar = 50 µm. B) Dome prepared with embedded fluorescent beads (left of dashed line) enabled visualization of further deformation of the edge. Different heights and transversal reconstruction show that the upper part of the dome moved further from the edge and that some cells are covered by the gel. Scale bars = 50 µm top, 10 µm, bottom. C) Confocal images showed that the structure at the edge was made of cells organized in a weaved manner. A difference in protein expression is clear between the tubular structure and the neighboring monolayer on glass. Scale bar = 25 µm. D) Corroborating this, a side view of the structure (following yellow line) showed a zigzag organization of the cells, and transverse cross section showed that the folding had closed the structure. Scale bar = 25 µm.

Interestingly, the lumen then appeared after a cell-autonomous coordinated process, due to the apical-basal polarization by laminin of these cells, facing each other inside the wrapped cord. Some reports indeed suggest that laminin polarization plays a crucial role in orienting epithelial and endothelial cells along an apical-basal axis, a necessary step for lumen formation ^19,46^. In order to confirm this, we evaluated the evolution of cell polarization between the start of stage 3 and the end of stage 4, as well as the maturation of tight junctions, the presence of a lumen and the remodeling of the ECM. Regarding podocalyxin localization, the front view of the structure in both stages revealed that it was two cells thick, which allowed the definition of a top and bottom layer. (Figure 9A). While podocalyxin was predominantly expressed at the top cell layer, with no signal detected in the bottom cells or between cell layers at the end of stage 3, in contrast, during stage 4, when the ECM had completely engulfed the two layers of the cords, podocalyxin was mostly found between the two layers (Figure 9B). This result confirmed our assumption that the ECM dome edge folding induced apical-basal polarization, with the apical side facing now the interior of the cord structure. This central inner apex of all cord cells was critical for subsequent lumen formation ^28^ as it was consistently found prior to the appearance of a lumen, which gradually emerged in what had now become a microvessel (Figure 3B and Figure 10).

**Figure 9.**
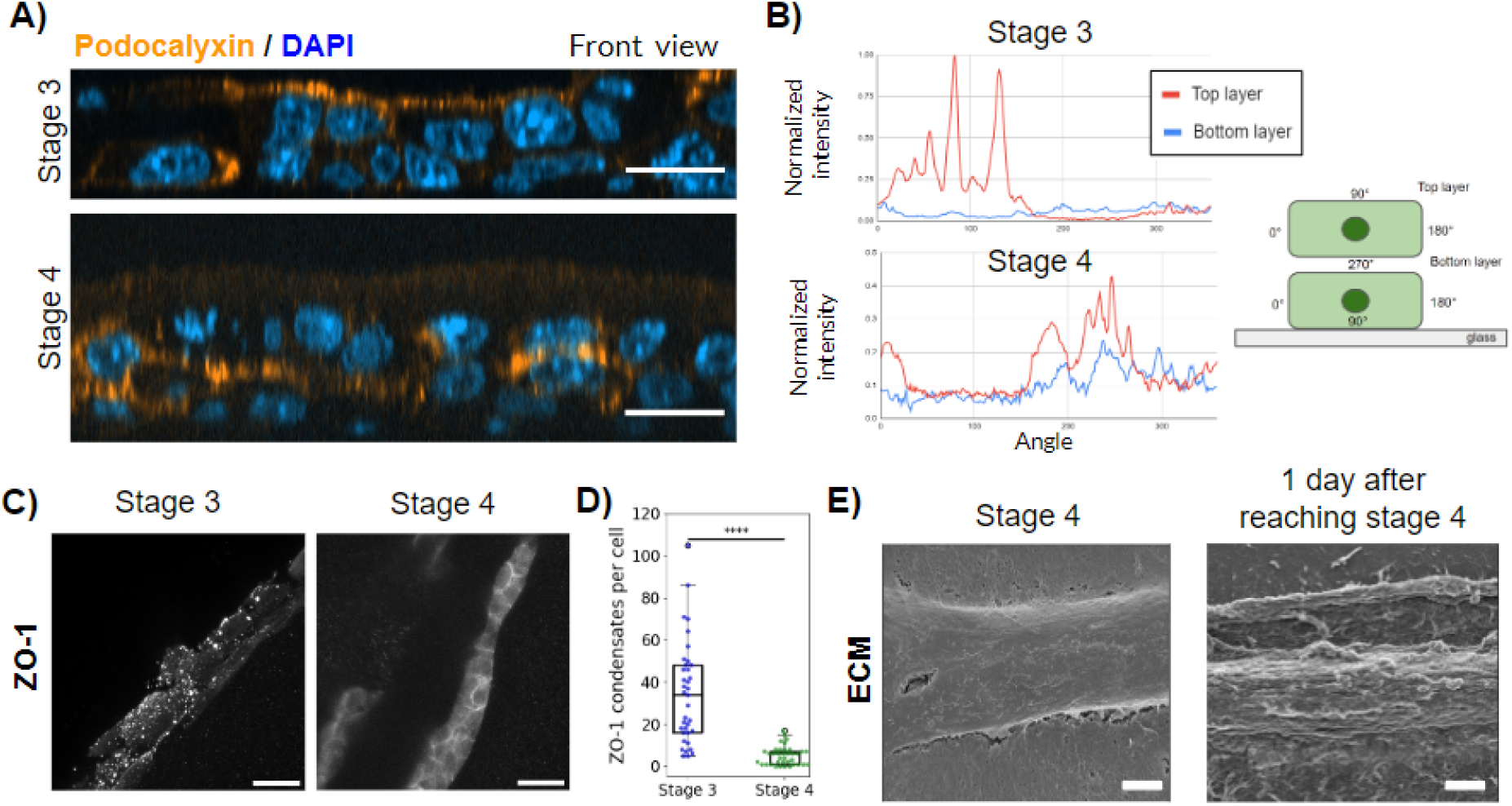
Complete folding of the GT dome promotes the maturation of a lumenized vascular structure. A) Front view micrographs of capillary-size structure stained for Podocalyxin (PDCLX) during stages 3 and 4. Scale bar = 15 µm B) Cell periphery intensity profiles were obtained for pairs of cells (top layer and bottom layer, as shown in the schematic). Quantified normalized intensity as a function of the angle shows a clear change in localization from the top part of the structure (angles 0°-180° in red) to in between layers (angles 180° - 360°, red and blue) can be seen. n = 2. C) Confocal micrographs show the expression of ZO-1 transfers from intercellular condensates in stage 3, to a cell-cell junction localization at stage 4. Scale bar = 15 µm. D) Number of ZO-1 condensates per cell in the structure decreases in stage 4 compared to stage 3. Data were analyzed using Mann-Whitney U test to determine significant differences between groups. (n=2, “^****^” P<0.0001). E) SEM micrographs showing the evolution of the outer part of the vascular structure after stage 4. A thin ECM layer reorganized and formed wavy bundles. Scale bar = 2 µm.

**Figure 10.**
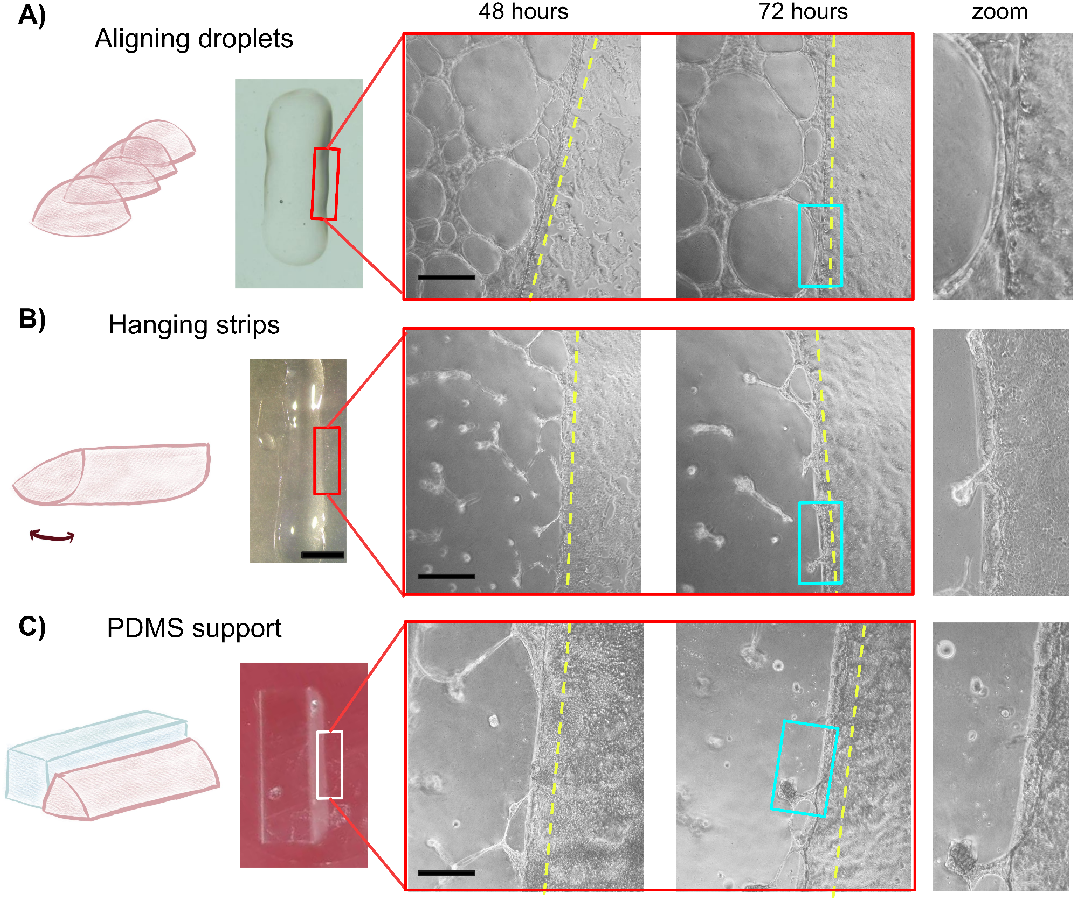
Straight-vessel guidance fabrication strategies. GelTrex deposits were engineered in three different ways for TSEC culture: A) aligned semi-coalescing domes, B) hanging strip, and C) alongside a PDMS scaffold support. After 48 hours, cords are visible at the edge of the architectures (yellow dashed lines). Typical vasculogenesis networks (A) or aggregates (B-C) are visible in the central regions. Scale bar = 500 µm. The right column shows zooms of the edge after 72 hours of culture. The characteristic ECM displacement of the folding step is similar to those of Figure 3 and Figure 6.

The final step of microvessel engineering at the end of stage 4 consisted in the progressive maturation of the lumenized vessels into functional vessels with stable cell-cell junctions and thin BM secretion. In our experiments, the organization of ZO-1 clearly changed from cytosolic condensates in stage 3-TSEC cells into a prominent localization at the cell-cell regions in stage 4 (Figure 9C and Figure 9D). This transition has been reported elsewhere and was associated with a response to an increased inner pressure building up during mouse embryo development ^47^. Finally, a slightly visible thickening of the structure was found to correspond to a secretion and organization of a BM into wavy bundles, aligned by the cells in the direction of the vessels (Figure 9E). This last step of microvessel maturation by BM secretion was confirmed by immunofluorescence assays, confirming that collagen IV and laminin were constituents of the vessels BM (Figure S6).

All these findings indicate that stage 4 is the lumenization and maturation stage, comparable to those reported in the development of vessels *in vivo*. It was achieved spontaneously by cells after the completion of the previous stages, engineered by a 3D ECM geometry, necessary to guide cells towards the formation of a microvessel. A summary of the morphogenetic process, along with its corresponding scheme, is presented in the supplementary material and especially in Figure S8.

### 3.4 The direction of the vascular structure can be controlled by the initial shape of the guiding edge

To further enhance our approach of controlled gelation process and soft ECM edge formation without the need for a non-physiological circular shape, we next explored other configurations that allowed vessel development in a different geometry. We first tested the creation of a straight ECM line with similar properties needed to form the microvessels, by arranging multiple small GelTrex domes in a row, with a short separation for the domes to rapidly coalesce before gelation. Unfortunately, this procedure left small, but visible, narrowings between consecutive domes (Figure 10A). To avoid this periodic narrowing, continuous lines were arranged by extruding small gel volumes manually, through a pipette tip (Figure 10B). We also employed a straight, fixed slab of polydimethylsiloxane (PDMS), treated with UV-ozone to turn its surface hydrophilic while placing the gel droplet. A widely used material in microfluidic and MPS applications, the PDMS block served here as a support against which the gel could be deposited while maintaining the outer edge shape interface between ECM and glass, as required for our microvessel formation (Figure 10C). After 24 hours, for all configurations, cells attached to the edge of the GelTrex architectures similarly to what had happened for the previous domes. After 48 hours, networks of cells were visible in the middle areas of the gel, connecting with the cord formed at the edge, recapitulating the organization shown in the domes in the central and peripheral regions. Finally, after 72 hours, cords appeared at the edge and showed the same behavior as observed in the domes, with stage 4 evident in the zoomed images of Figure 10. This confirmed that altering the geometry of the GelTrex deposit does not hinder the self-organized vessel morphogenesis, as long the folding-permissive softness of the gel was maintained, as it is essential to ensure the proper verticality of the edge, avoiding a flattening on the outside.

## 4. Discussion

Vascularized MPS are essential to better understand the mechanisms involved in disease development and maintenance of homeostasis. However, in many existing *in vitro* models, vessels embedded in a 3D matrix often fail to imitate the architectures of capillaries surrounded by little or no basement membrane. This limitation hinders the faithful recapitulation of many tissue-vessel interactions, particularly those crucial for discontinuous microvessels not enveloped by ECM proteins or supported by pericytes or fibroblasts. Here, we demonstrate an alternative approach to generate non-embedded *in vitro* microvessel models, specially tailored for liver sinusoid. Our straightforward microvessel fabrication method allowed us to successfully guide LSECs to autonomously construct an *in vivo*-like morphogenetic process, resulting in a progressive microvessel formation with liver sinusoid characteristics. This approach holds promise for engineering MPS constructs that more accurately mimic physiological conditions.

First, scattered ECs were guided to autonomously form a mature microvessel with a preferential direction, observed in both murine and human LSEC cell lines. Notably, this distinct process does not rely on the support of prefabricated 3D tubular scaffolds ^48^, interstitial flow inside a hydrogel ^8,49^, or stromal cells ^50^. We identified the critical parameters of the microenvironment essential for engineering a not-clamped lumenized structure. By leveraging the composition and anisotropic micropatterning of laminin, a protein abundant in early development of vessels, ECs self-organize sequentially, mirroring the morphogenetic processes of vasculogenesis observed *in vivo* ^27^. Cell migration and proliferation were guided by this pattern that was carefully shaped via the precise control of the Marangoni flow. They resulted in the formation of a 1-cell (1D) train followed by a multicellular (2D) cord and a (3D) lumenized microvessel tube. Under mechanical tension and shear probably due to the localized proliferative and migrational regimes at play, cells then folded a thin outer portion of the gel and wrapped the cord with ECM. The enriched outer presence of laminin around the wrapped cord facilitated the polarization of cord cells, leading to the opening of a lumen in their interior. This process ultimately resulted in the formation of a vessel that autonomously remodeled its own ECM and size.

Flat basement membrane-derived substrates are commonly used *in vitro* for self-structuring, often forming endothelial cords ^15,16^. Recently, the interplay between the physical plasticity of laminin (as opposed to collagen’s elasticity) and the contractility of cells has been highlighted as a crucial factor for long-range cell-cell mechanical communication underlying such cord formation ^23,24^. As a result, cells organize into complex network architectures that resemble the primary capillary plexus of vascular development, despite having a relatively flat, randomly oriented, and the absence of lumen in these multicellular structures. This limitation can potentially be circumvented with an additional interstitial flow or ECM overlay onto the structures, to provide a polarization signal that opens a few localized hollow regions thanks to the resulting shear stress ^6,51^. Here, following developmental biology insights of tissue-wide folding processes, we proposed that a long multicellular cord could be engineered along a laminin micropattern, to anisotropically direct and orient a single 2D cord thanks to the localized shear stress caused by the proliferation and migration patterns that were restricted to the preferential direction set by the pattern. The use of a fixed ECM pattern on the cell basal side, typically obtained by microcontact printing ^39^ is indeed insufficient for lumenization, due to the absence of apical-basal polarization. To address this limitation, we adapted a controlled Marangoni flow process inspired by the work of Nerger *et al*. from Celeste Nelson’s group ^29^ to a GelTrex gel dome. Although laminin is not a fibrous protein, we leveraged the infamous “coffee stain effect” to achieve two important cues simultaneously. First, a thin laminin micropattern condensed at the edge of the dome, facilitating anisotropic micropatterning of laminin along the dome edge. Then, the vertical (3D) edge introduced a local ECM wall perpendicular to the glass on which the cell cord axis aligned. It acted as a physical cue that facilitated ECM folding upon the cord, polarizes the cells in the apical-basal axis and completes the necessary developmental steps for lumen formation, as discussed below.

We observed that the patterned ECM triggered a first “approach and adhesion” stage for some cells initially located nearby, on the surrounding glass. Our results consistently showed an attraction distance threshold of approximately 50 µm, suggesting that this first stage was governed either by chemotaxis, a migration regime induced by a gradient of soluble factors, or by haptotaxis, a migration regime induced by an ECM gradient. Both motility modes have been reported to drive EC morphogenesis ^52^. Notably, the commercial ECM (GelTrex) used in our study contains various growth factors, including VEGF, which likely attract nearby cells exposed to a diffusive gradient. In addition, cells adhering to soft gels can extend the release of more molecules embedded in the gel over time by exerting pressure on the soft and porous ECM ^53^. This makes soluble factors more accessible through diffusion, particularly in close proximity to contractile cells such as LSECs ^43^. Upon adhesion, cells proliferated and migrated exclusively along the perimeter of the dome, forming a persistent, anisotropic, 1-cell-thick train that rapidly lined the edge. We propose that the anisotropic edge of the dome and its composition were responsible for the collective cell polarization and orientation observed in this stage, thus preventing the random orientation of cord segments typically observed in conventional 2D assays. When a spread layer of GelTrex was tested, the protein pattern at the edge was lost, resulting in the formation of short, disconnected segments instead of the unique cord. We believe the specific geometry of the edge was not sufficient to promote this unique cord organization. Fluid and solid shear conditions were shown to influence endothelial morphogenesis in other studies ^54^. The mechanical properties of the GelTrex edge combined with solid shear along the thin laminin micropatterns provided an ideal ground for robust cell alignment, as reported for cell cord formation ^24^. Indeed, when testing a dome of collagen I instead of GelTrex, ECs failed to form the train and instead covered the dome with a monolayer. Furthermore, when laminin is absent, ECs tend to aggregate into continuous sheets ^20^. Fibrous collagen IV, also present in our ECM dome, has been reported to direct migration in aortic ECs ^55^, while laminin plays a pivotal role in polarizing other cell types and defining their direction of migration ^19^.

Subsequently, the organization transitioned from a single peripheral train to a multicellular cord, which would later construct the microvessel. We hypothesize that a gradual shift in cell polarization, driven by shear, was at play, akin to the observations in retinal capillaries morphogenesis. It has been shown that laminin 511 strengthens cell - cell junctions, aiding ECs to resist shear stress ^56^. In this context, microvessels emerge and organize through different polarization states, responding to competing signals from VEGF and shear stress sensed at focal adhesion. Depending on the direction of polarization induced by these signals, they instruct either the building of new structures or their remodeling ^57^. In our study, cells first formed a single peripheral train along the whole perimeter, seemingly driven by preferential division and migration. Only once the entire edge was covered did the proliferation direction change in some locations where cells started to divide perpendicularly and climb up the edge, to form a multicellular cord that would then bend and fold the gel. The change in direction observed could be mediated by Planar Cell Polarization (PCP), a polarity axis and signaling pathway that is critical in vascular morphogenesis ^58^. Studies have indicated that PAR-3 regulates PCP and can be modulated by fluid flow ^59^. Additionally, flow-associated shear has been shown to remodel the laminin-integrin network ^60^. The influence of shear stress on endothelial cell PCP is well-documented, with evidence suggesting reversible effects upon modification of shear direction ^61^. In this scenario, the migration and proliferation of cells on the high-density laminin pattern may have promoted shear stress signaling, as observed in similar studies ^56^. However, further investigation is needed to explore the role of PCP in this context.

Interestingly, this multicellular cell cord transitioned from 2D to 3D architectures through folding. We disregarded local apical constriction as the underlying mechanism for tissue deformation, as we did not observe the characteristic cellular organization of F-actin which is typically associated with this process on a wide scale deformation of tissues in development. Instead, a long-range 2-step pulling and invaginating process was identified, accompanied by a laminin depletion at the basal side of the hinge cell. Similar events have been observed in Drosophila development studies, where localized ECM depletion was noted during tissue-wide folding in wing imaginal discs ^45^. Moreover, during the invagination of epithelial cells forming the cephalic furrow, recent findings revealed that infolding serves as a mechanical buffer, releasing energy that accumulated due to tissue growth in a confined volume ^62^. Notably, this buckling occurs between two groups of leading cells, tensing the central adjacent region before invagination. This morphogenetic process occurring at the edge of the soft dome strikingly resembles that of the mechanical selection of leader cells by polarization through a lateral tensile load in collective migration, as first described by Spatz group in 2018 at the edge of epithelial monolayers ^63^. This invagination guided by the cell anisotropic alignment thus involves traction patterns that propagate along several cell diameters and determine when and where invagination occurs. In our study, gel folding indeed always occurs between two cell clusters that look like mitotic groups or leader cells, polarizing orthogonally to the edge and tube in formation. By initially imposing a lateral strain to this confined region, the mechanical instability linked to stretching resembles a shear-driven plane buckling, caused by a tensile load, as described for non biological materials ^64^. This phenomenon, stunningly similar to the patterns found in our experiments, occurs when a soft sheet of material suffers an out of plane deformation upon extended longitudinal tension ^65^, resulting in out-of-plane longitudinal wrinkles or infolds ^66^. The direction, length and deformation threshold needed to initiate these deformations are defined by the interplay between external stress and orientation of the superficial or inner substructure of the material ^67^. In our study, we engineered a lateral invagination process where cells’ pulling on the ECM pattern, our anisotropically pre-oriented structured material, generate the necessary shear to trigger a folding. A certain length of persistence seemed necessary for the folding to appear and wrap full cord segments, which was evidenced by a periodic repetition of invaginated segments of approximately 300 µm in all dome experiments (regardless their original size), that eventually connected to form the full circle vessel. Interestingly, this is twice the length of separation found by Vishwakarma et al. ^63^ between their leader cell groups. In their work, they attributed this distance to the distribution of force correlation length, which, in their experiments, is provided by a bulk layer of cells behind the edge sitting on a stiff substrate, probably limiting the force correlation length. In our case, cells are organized in a 2 to 3-cell thick line, pulling on a very soft ECM along a thin, dense laminin edge, possibly enhancing the effect and widening the force landscape to a higher correlation length. Consolidating this hypothesis, as mentioned earlier, we found that when the external monolayer grown on glass managed to invade the edge and prevented proper cell alignment and cord formation pre-invagination, this important folding step did not occur.

No long-term, sustained tension was observed in this case. Verifying this assumption would be interesting to fully explain the folding step in our vessel engineering process, but it falls outside the scope of this present study, and we found this process to be extremely robust.

Although the tension from these deformations can lead to spatial signaling that modulates vascular morphogenesis ^68^, the folding also allowed a complete self-wrapping of the multicellular cord by a thin layer of GelTrex. ECM plays a crucial role in regulating lumen formation by establishing endothelial cell polarity ^69^. In particular, laminin orients the apical pole of cells through Rac1 and ß1 integrins ^18^ via PAR-3 ^59^. Notably, progression toward lumen formation in our model mirrored maturation events observed in *in vivo* lumenogenesis ^28^. First, the polarized trafficking of podocalyxin was observed towards the center of the multicellular cord formed after folding, establishing this protein as the apical surface from which the lumen emerged ^70^. Then, tight junction protein ZO-1 and F-actin were recruited at this region, ensuring lumen maintenance in spite of a probable pressure build up ^71^. Finally, the multicellular cord, initially enveloped in a very thin ECM blanket after folding, was surrounded by an organized structure of laminin and collagen IV that resembled a thin basement membrane from a mature vessel ^72^. Electron microscopy revealed a striking evolution between the end of the folding process and a subsequent stage one day later. Thin, short, wavy fibrous bundles organized and aligned with the direction of the underlying vessel, forming a supramolecular network characteristic of a BM ^73^, without a noticeable overall increase in vessel thickness.

In conclusion, the anisotropic formation of a thin cell cord was enabled by the engineering of a 3D ECM edge boundary. It favored the cell-driven folding of a soft, laminin-rich hydrogel that polarized the endothelial cell to gradually open a lumen and mature the junctions and basement membrane. As recently mentioned in a study, the generation of anisotropy in geometrical elements allows shear-align cells ^74^, and it is crucial in endothelial vasculogenesis assays to form stable microvessels ^23,24^. We have described each stage of this process with precision and demonstrated how to engineer them with minimal external (and only initial) intervention. To the best of our knowledge, the orchestrated process—involving the anisotropic edge, formation of an EC cord, a wake-like folding and further maturation of the cord—has not been previously evidenced for ECs. While we acknowledge that the cell source could be modified to create a more accurate sinusoid model, we anticipate promising outcomes from our constructed vessels, with future work focused on their characterization as tissue-specific capillaries and their manipulation and integration into MPS to facilitate perfusion of such a delicate and bare microvessel and provide more physiologically relevant tissue-vessel interactions.

## Acknowledgements

A.X.M.R. conducted this study to fulfill the requirements of Programa de Doctorado en Ciencias Biomédicas, UNAM, and received a doctoral scholarship from Consejo Nacional de Humanidades, Ciencias y Tecnologías (1004395). The authors are thankful to Michaël Trichet and Alexis Canette from the IBPS electron microscopy core facility and France Lam from the IBPS microscopy core facility (Sorbonne-Université, CNRS) for their support in imaging. Special thanks to Pierre-Emmanuel Rautou for kindly providing the TSEC cell line and to him and his team for engaging in discussions. We also thank Marina Macías-Silva for providing the TMNK-1 cell line. We extend our gratitude to Benoit Ladoux, René-Marc Mège and Sushil Dubey for providing access to their facility and for continuous support in this project. Lastly, our greatest gratitude goes to Michel Labouesse and his team for hosting us. The team was funded by the i-Bio initiative and supported by a DIM ELICIT grant from Région Ile-de-France.

## Competing Interest Statement

The authors declare no competing interest.

## Author Contributions

A.X.M.-R and M.H. designed research; A.X.M.-R, B.N.-R, W.X. and M.H performed research; A.X.M.-R analyzed data; A.X.M.-R and M.H. wrote the paper, A.X.M.-R, B.N.-R, W.X. and M.H reviewed the paper; A.X.M.-R, W.X. and M.H edited the paper.

## Notes

### Competing Interest Statement

The authors have declared no competing interest.

### Summary of Updates

Improved writing and explanations after version #1

