## Supplementary information for "Microvascular engineering for the development of a non-embedded liver sinusoid with a lumen: when endothelial cells do not lose their edge"

**This PDF file includes:**

Legends for Movies S1 to S4
Supporting text
Figures S1 to S8

---

Legends for Movies S1 to S4

Movie S1 (separate file). Phase contrast microscopy movie of TSEC organizing during stage 1.
Movie S2 (separate file). Phase contrast microscopy movie of TSEC organizing during stage 2.
Movie S3 (separate file). Phase contrast microscopy movie of TSEC organizing during stage 3.
Movie S4 (separate file). Phase contrast microscopy movie of TSEC organizing during stage 4.

Supplementary Text

### Deciphering optimal substrate conditions for guiding LSEC organization.

On a technical note, optimal conditions for the fabrication of GT domes were determined through a series of experiments. We observed that careful handling of the domes is absolutely essential for reproducibility and desired outcome. Experimental outcome was highly sensitive to the initial conditions as discrepancies in cell seeding density, dome contact angle, GelTrex stiffness and culture medium components have led to rapid covering of a monolayer or numerous sprouts appearing at the edge, below the gel; both conditions impeding the morphogenetic steps necessary to form the vessel.

For instance, allowing the gel to dry at low humidity or for a longer time in the incubator, and leaving it without culture medium after gelation are unwanted as they all changed the dome properties, increasing stiffening and disturbing the final organization of the cells, preventing cord formation and maturation and promoting the development of a monolayer (Figure S2). On the other hand, manually spreading the droplets resulted in a completely different cell organization outcome (Figure S3). We hypothesized that this was due to the disruption of a cell-size laminin pattern at the beginning of the experiment. The pattern resulted less pronounced at the edge of the structure, with more diffuse and uniform distributions that were larger than a cell size, confirmed by profiles from the immunofluorescence. Interestingly, this led to an interrupted, piecewise tube-like organization, different from that found on intact domes (shown in main text, Figure 3A). This suggests that, similarly to what was found by Nelson's group using collagen I, the organization of non-fibrous, laminin-rich ECM with Marangoni flow also controls adherent cell future patterning.

### Simple visualization of proposed folding process.

The observed mechanism of the folding following lateral strains that leads to the encapsulation of cells by a laminin-rich ECM is not easy to understand intuitively. We kindly suggest the readers to observe what we believe is the closest phenomenon with a simple test. Take a thin piece of soft cloth (e.g. a microfiber cleaning cloth for glasses), grasp it firmly with your two hands between the index and thumb and gently pull it apart. A transverse buckling and folding appears immediately. Intuitively, the greater the distance between the two pulling anchors, the less shear strain required to obtain the buckling (visible as a wrinkling in the piece of soft fabric). This last observation may help explain why there seems to be a minimal cord length needed for the folding process to occur, as regulated by gel edge softness (hence buckling resistance). More experiments employing traction force microscopy with a greater spatiotemporal resolution and nanoindentation to obtain the exact elasticity of the edge will be necessary to try and determine the forces at play here.

### Summary of the morphogenetic process

The Figure S8 recapitulates the gradual morphogenetic vessel formation described in the manuscript, detailing the stepwise morphogenetic process followed by cells and that was evidenced by our experiments. LSEC spontaneously formed a lumenized capillary-sized vessel. This was achieved by murine and human cells, without external assistance from any mural or stromal cells. The only exogenous support was the presence of an edge of a GelTrex soft matrix, necessary to initiate the whole process which is then completely cell autonomous. This boundary condition was ensured by the Marangoni flow-driven gelation of a GelTrex droplet: it created an ECM density gradient culminating in a thin pattern of concentrated laminin that strictly follows the edge of the dome. This pattern, of one cell-size width, attracted the migration of the endothelial cells from a 50 µm vicinity. Upon adhesion to this ring-like pattern, cells followed this trail: anisotropic migration and proliferation guided a gradual organization of sparse, individual cells into a 1-cell ring-like cord (1D). Upon covering the whole edge, proliferation locally changed to form a multicellular cord (2D) which spread planarly over 2-3 cell widths. Local lateral deformations of the gel provoked a vertical folding process, reminiscent of a surf wave, around a line of hinge cells. The progression of the fold is similar to that of a wave, breaking locally by plunging and extending sideways. Part of the dome advanced towards the gel-glass interface (corresponding shore following the wave analogy). The plunging part curled and wrapped and engulfed the whole multicellular cord, which became an ECM-covered, capillary-sized tube. An apical-basal axis then established, evidenced by podocalyxin localization and the structure entered a lumenogenesis process. It correlated with ZO-1 tight junction maturation and F-actin recruitment in the apical zone and ended up in a microvessel tube with a lumen, running all around the dome. This vessel then slowly matured, as evidenced by the thin BM formation and organization in the direction of the vessel.

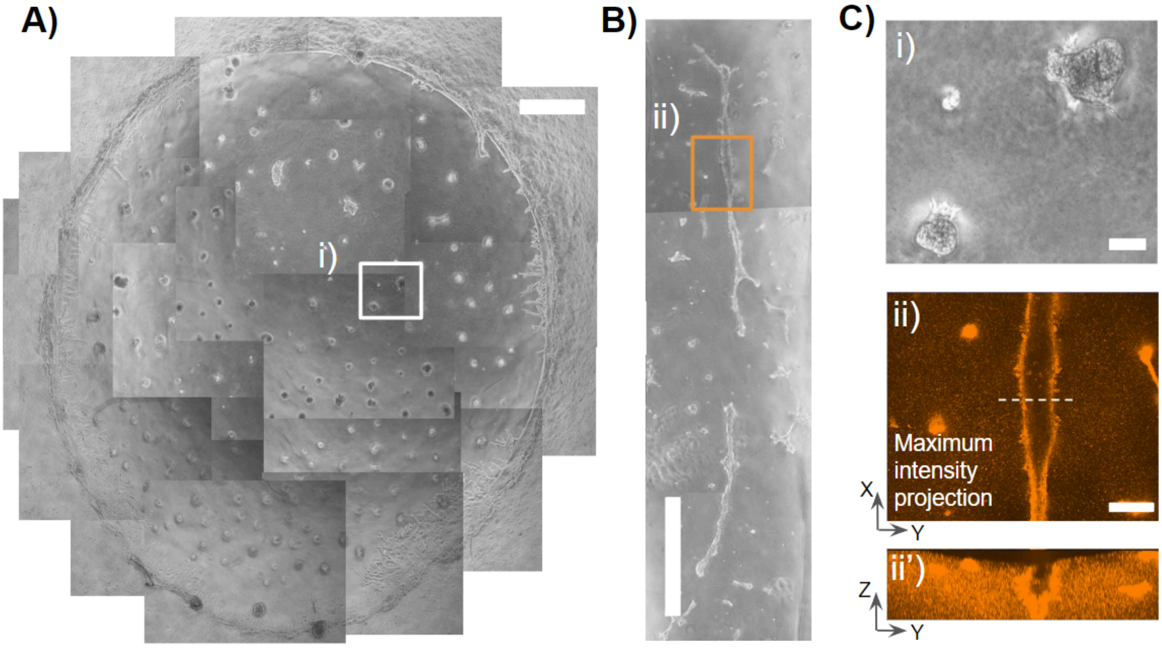


**Supplementary figure 1. Deposited GT guided ECs either to spheres or cords depending on its geometry** A) Top-view brightfield image of TSECs cultured for 72h on an isotropic laminin condition. Center networks eventually detached from their ends and tangled up into 3D aggregates of various sizes, scale bar = 1 mm. B) Top-view brightfield image of TSECs cultured for 72h on a 2D anisotropic laminin condition, scale bar = 1 mm. Gel was placed between two PDMS slabs and cells formed long cords C) Zoomed regions from both conditions: i) two aggregates from the center of the isotropic laminin condition, scale bar = 100 µm. ii) Maximum intensity projection of structure stained with WGA. Scale bar = 200 µm. In the cross section (ii´) it is observed that the structure is semi-closed, not completing the folding nor forming a tube.

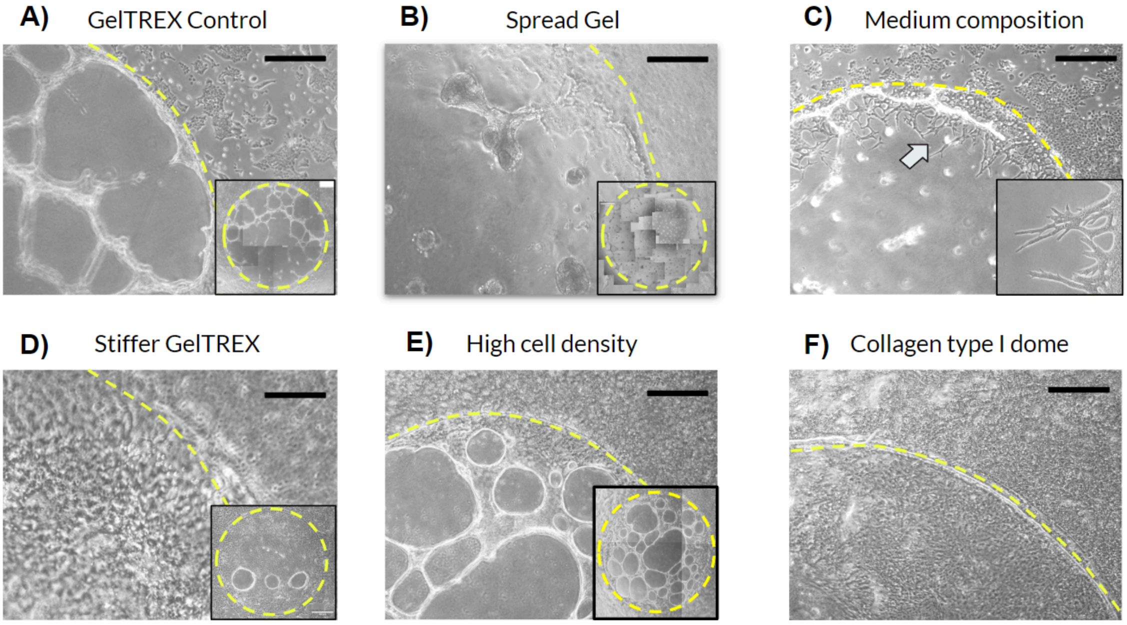


**Supplementary figure 2. Gelation conditions (gel composition, architecture and environmental cues) directly impact the cell organization.** Comparison of bright field micrographs showing TSEC organization after 72 hours of cell culture on different substrates. Yellow dashed lines indicate the border of the gel. Scale bar = 500 µm. A) Geltrex dome control where a cord organization is recapitulated at the edge as well as a network in the middle part. Inset shows the complete dome observed. B) Spread Geltrex droplet prior gelation. The network in the middle is lost at 72 hours and the edge is composed of short segments rather than a complete structure as in the control condition. Shown in inset the whole dome. C) The use of medium without any culture growth factors promoted the rapid emergence of sprouts from cells at the edge towards the dome. Inset shows sprouts which seem to be directed towards the cell aggregates at the center. D) Failure in controlling the gelation/evaporation ratio promoted an increase in gel drying which we hypothesize increases the stiffness of the material. In that case the cells formed a monolayer that covered most of the dome as seen in the insert. E) When the density of cultured cells was greater than that established 196 cell/mm2, it was observed that on the dome there were areas or islands that more closely mimicked the monolayer morphology than the networks observed in the control condition. These monolayer islands increased in size over time. F) To verify that it was not just the edge that provided the organization, we manufactured a type I collagen dome. In this case, it was observed that the cells formed a monolayer that covered the entire surface of the dome.

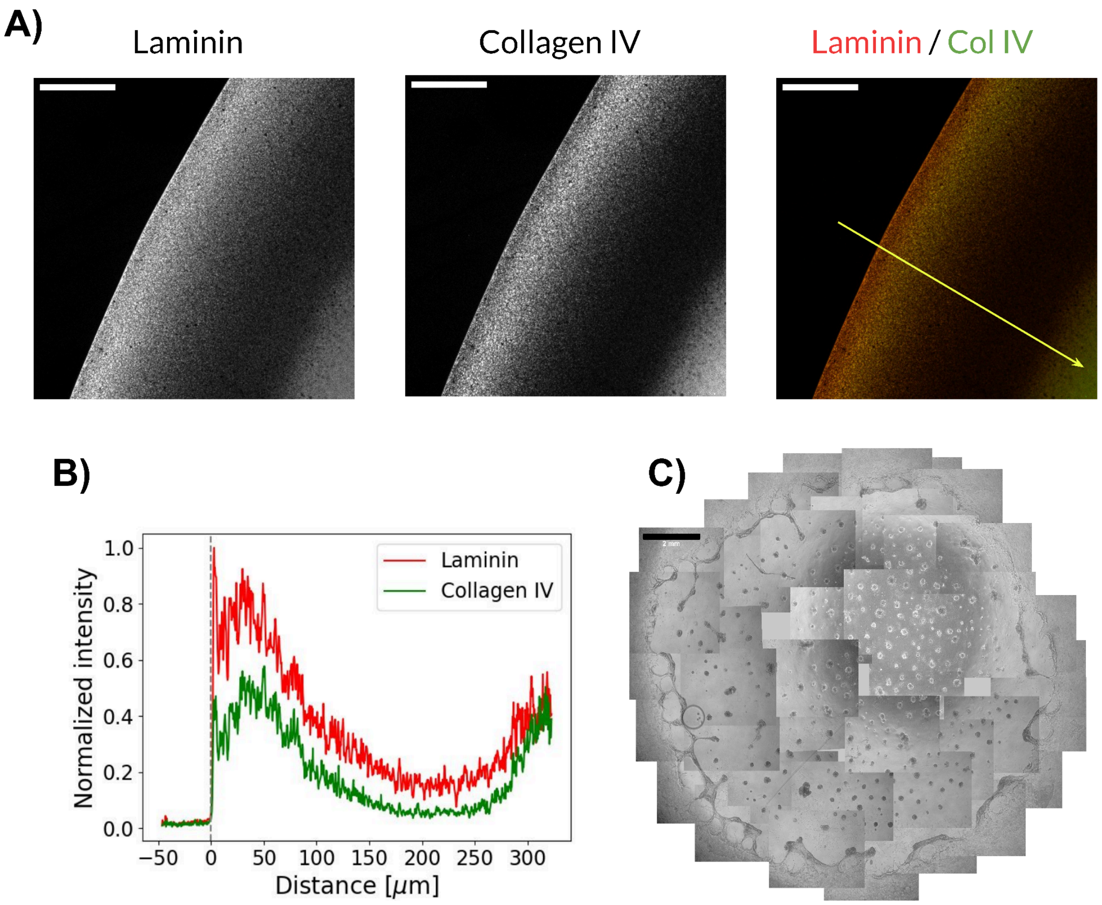


**Supplementary figure 3. Spreading the Geltrex droplet prior to gelation yields an homogenic deposit of laminin and an incomplete cell cord.** A) Confocal Z projections spread droplets stained for Laminin and Collagen IV showing an homogenous deposit of protein. Scale bar = 100 µm. B) Profiles of lines going transversally and inwards through the spread droplet (as shown with the yellow arrow) were obtained both for laminin and collagen IV. We show the average profile for both stainings with a distance of 0 µm corresponding to the edge of the dome. n = 4 per group C) Top-view brightfield image of TSECs cultured on a Geltrex spreaded dome at 72h. Scale bar = 2 mm. At the edge we observed short disconnected segments of cords rather than a unique cord around the edge.

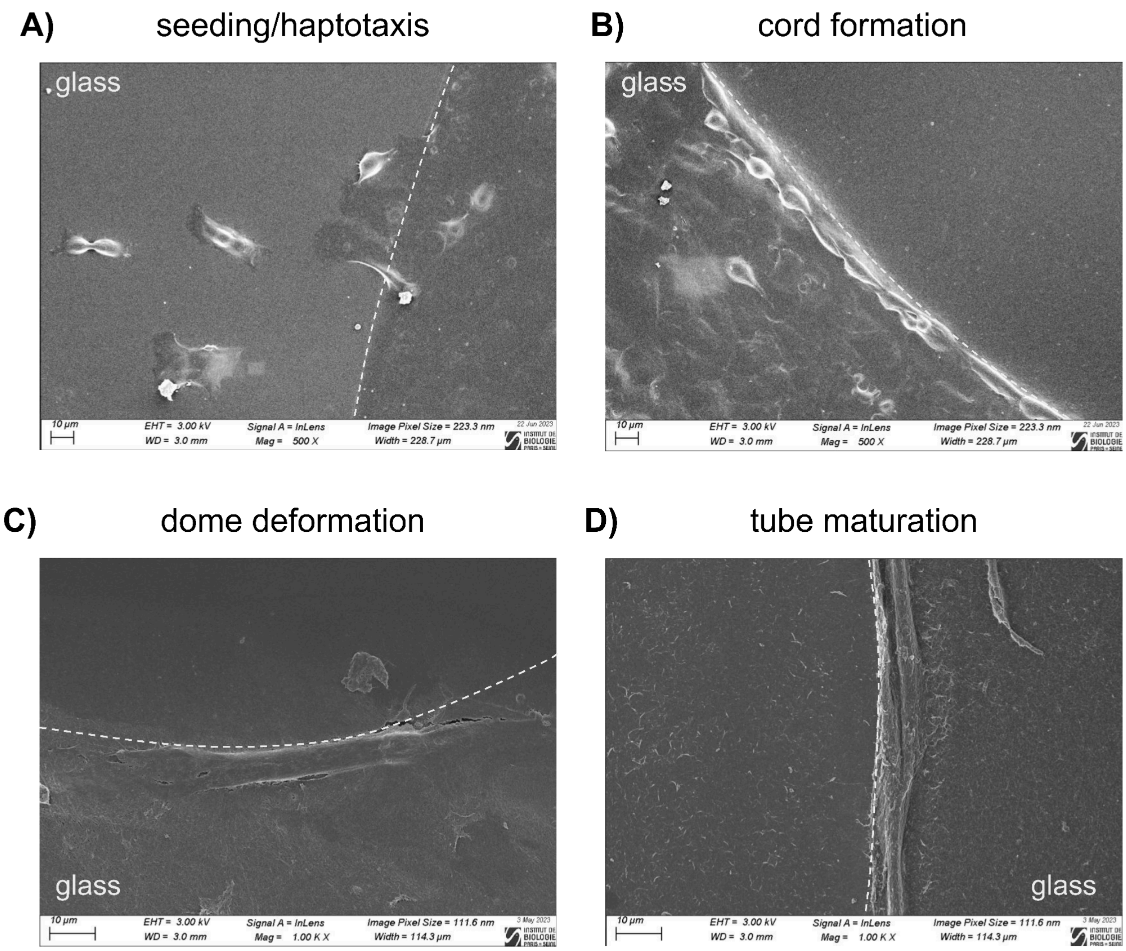


**Supplementary figure 4. Endothelial cells follow specific morphogenetic steps to form a mature vessel, showcased by SEM micrographs.** A) Beginning of the organization with individual cells migrating towards the GT dome. B) Cord formation by cells proliferating and migrating on the edge. Cells are not densely packed in a cord yet. C) A thicker structure can be seen at the edge of the dome, in this representative picture the folding has only covered a short distance. D) Finally a structure covering the whole edge can be observed also with a topography that suggests the deposition and organization of ECM proteins. Scale bars = 10 µm.

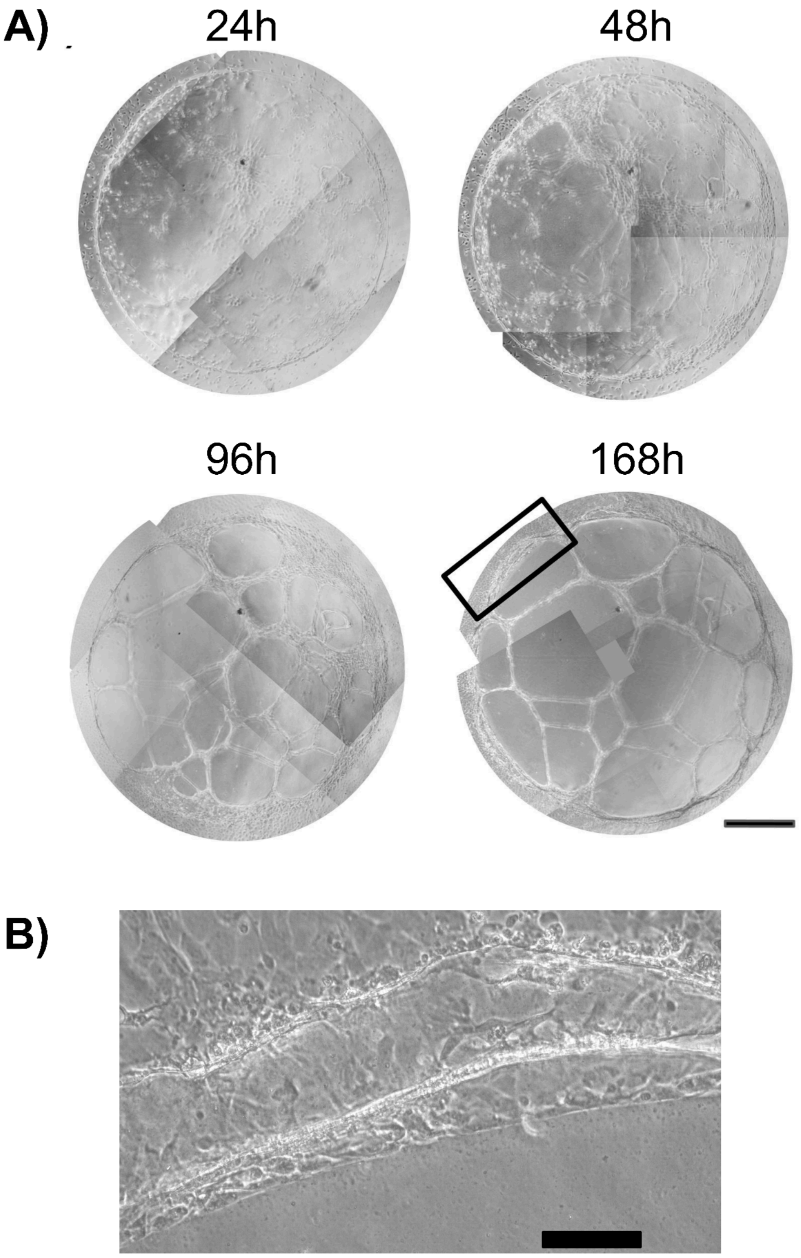


**Supplementary figure 5. Immortalized human LSECs are also organized into a tubular structure at the edge of a gelTrex dome.** A) Temporal evolution of TMNK-1 organization on gelTrex domes. We can observe the formation of networks in the central part that connect with the edge. Scale bar = 500 µm B) Zoom of zone of interest at the edge of the dome. We can observe the characteristic edge of the GelTrex, covered by a folding and wrapping event. Scale bar = 100 µm

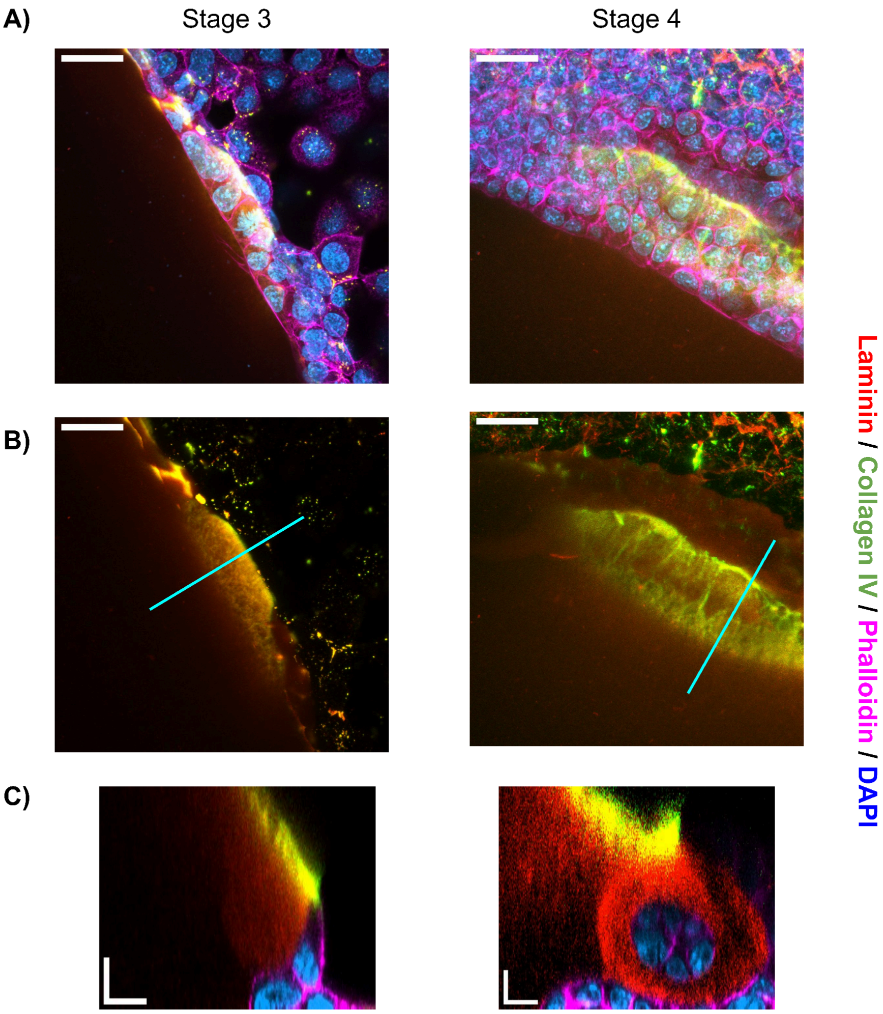


**Supplementary figure 6. Evolution in the organization shows the wrapping of the vascular structure with extracellular matrix proteins and in particular the colocalization of laminin and collagen IV.** A) Z projection of staining against laminin, collagen IV and labeling with phalloidin and dapi. It is shown that the matrix covers the cells and that this front advances as the stages progress. Scale bar =25 µm. B) Z projection of the laminin and collagen IV label showing colocalization in both stages and a progression of laminin covering the cells in the monolayer in stage 4. Scale bar = 25 µm. C) Transversal cross-section comparison of stage 3 and stage 4. Stage 3 shows the beginning of folding with the collagen IV signal in the superficial part of the dome whereas stage 4 shows that the folding has ended up wrapping the structure and that collagen IV remains in the superficial part of the dome. Scale bar = 10 µm

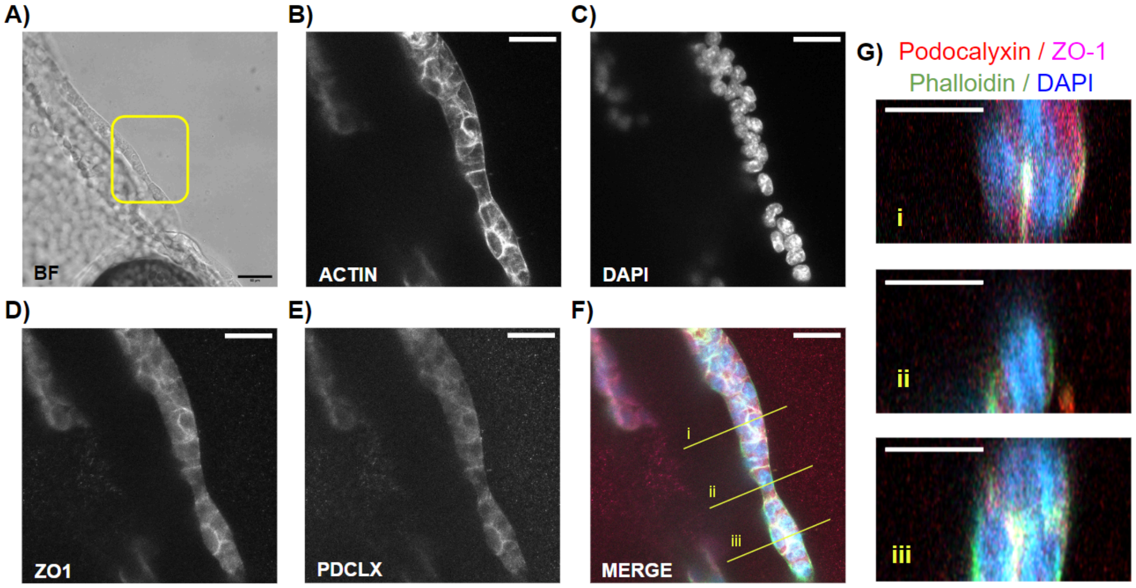


**Supplementary figure 7. Detail of the piecewise vessel formation around the dome**. A) Bright field micrograph of a vessel forming around the dome's edge, separating the monolayer (left) from the dome (right) Scale bar = 50 µm. Yellow square shows a zoomed region where staining of actin (B), nuclei (C), and against ZO-1 (D) and podocalyxin (E) was performed. Merge is presented in F) Scale bar = 25 µm . G) Transversal cross-section of the structure shown in F). It is observed that i) and iii) are two regions with proteins located in the internal part of the structure corresponding to a maturing vessel, connecting with a less mature region (ii). Scale bar = 15 µm.

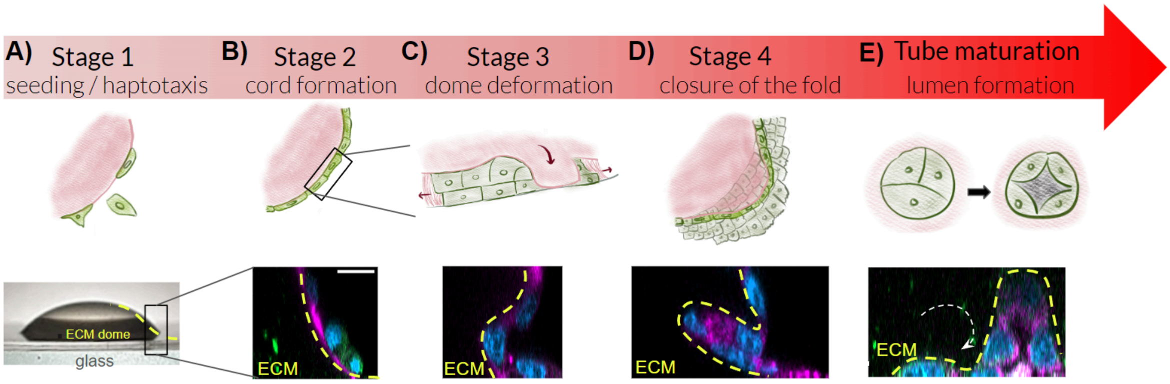


**Supplementary figure 8. GelTrex dome promotes a morphogenetic process leading to a lumenized vascular structure.** A) Initiation and 1D train: Marangoni flow-driven gelation of a GelTrex dome concentrated a ring-like laminin pattern to which cells were attracted and adhered preferentially to cover a 1-cell wide trail. B) 2D cord: tangential-then-upwards migration and proliferation transform the train into a flat multicellular cord over the surface of the edge. C) Folding initiation: in many locations, lateral tangential deformations fold the ECM gel. D) 3D wrapped cord: the gel progresses outwards and enrolls the whole cord; it is immediately followed by apico-basal polarization, with apical side inwards. E) Lumen opening: after polarization, a lumen is open. After some time, while lumen is preserved, a basement membrane is matured around the microvessels
